# *Map3k2*-Regulated Intestinal Stromal Cells (MRISC) Define a Distinct Sub-cryptic Stem Cell Niche for Damage Induced Wnt Agonist R-spondin1 Production

**DOI:** 10.1101/723221

**Authors:** Ningbo Wu, Hongxiang Sun, Xiaoyun Zhao, Lei Chen, Yuanyuan Qi, Yuheng Han, Xianan Liu, Caixia Gao, Qun Wang, Lingjuan He, Xiaoyin Niu, Zhiduo Liu, Hua-Bing Li, Yi Arial Zeng, Manolis Roulis, Dou Liu, Zhengfeng Yang, Bin Zhou, Richard A. Flavell, Bing Su

**Affiliations:** Shanghai Institute of Immunology, Department of Immunology and Microbiology, and the Minister of Education Key Laboratory of Cell Death and Differentiation, Shanghai JiaoTong University School of Medicine, Shanghai 200025, China; Yale Institute for Immune Metabolism, Shanghai JiaoTong University School of Medicine, Shanghai 200025, China; The State Key Laboratory of Cell Biology, CAS Center for Excellence in Molecular Cell Science, Shanghai Institute of Biochemistry and Cell Biology, Chinese Academy of Sciences, University of Chinese Academy of Sciences, Shanghai 200031, China; Department of Immunobiology, Howard Hughes Medical Institute, Yale University School of Medicine, New Haven, CT 06520, USA

## Abstract

Intestinal stem cell propagation and differentiation are essential for rapid repair of tissue damage in the gut. While intestinal stromal cells were recently identified as key mediators of this process, the cellular and molecular mechanisms by which this diverse population induces tissue repair remains poorly understood. Here we show that *Map3k2* has a colon stromal cell specific function critically required for maintenance of *Lgr5^+^* intestinal stem cells and protection against acute intestinal damage. This *Map3k2*-specific function is mediated by enhancing Wnt agonist R-spondin1 production. We further reveal a unique novel cell population, named *Map3k2*-regulated intestinal stromal cells (MRISC), as the primary cellular source of R-spondin1 following intestinal injury. Together, our data identify a novel intestinal stem cell niche organized by MRISC, which specifically dependent on the *Map3k2*-signaling pathway to augment the production of Wnt agonist R-spondin1 and promote regeneration of the acutely damaged intestine.

**Highlights:**

1. *Map3k2* protects mice from DSS-induced colitis by promoting intestinal stem cell regeneration.
2. *Map3k2*-MAPK pathway cross-talks with Wnt signaling pathway via upregulation of R-spondin1.
3. *<u>M</u>ap3k2*-*<u>R</u>*egulated *<u>I</u>*ntestinal *<u>S</u>*tromal *<u>C</u>*ells (MRISC) marked by co-expression of CD90, CD34 and CD81 defines a novel colonic stem cell niche.

## Introduction

Stem cell proliferation and differentiation are critical for rapid repair of intestinal injury and maintenance of host health (Nanki et al., 2018). Intestinal stem cells (ISC) are thought to rely on specific signals and growth factors produced by the local niche that support their repair functions and self-renewal in the crypts (Aoki et al., 2016; Degirmenci et al., 2018; Greicius et al., 2018; Kabiri et al., 2014; Sato et al., 2011; Shoshkes-Carmel et al., 2018). In particular, diverse populations of intestinal mesenchymal stromal cells (IMSC) appear to play key roles in ISC niche maintenance via sensing of factors including FGF, TGF-β, Hedgehog, and PDGF (Koch, 2017; Roulis and Flavell, 2016; Valenta et al., 2016), but the specific IMSC subsets and molecular mediators involved are unknown.

Several studies have now suggested that CD90^+^ IMSC may be crucial regulators of gut homeostasis (Huynh et al., 2016; Karpus et al., 2019; Kinchen et al., 2018; Powell et al., 2011; Roulis et al., 2014). However, recent efforts to understand the heterogeneity and function of ISMC using single-cell RNA sequencing (scRNA-seq) and lineage tracing techniques have revealed the existence of multiple functionally distinct subsets (Kinchen et al., 2018; Nanki et al., 2018), many of which produce soluble factors and cytokines likely capable of modulating gut homeostasis and epithelial integrity (Han et al., 2018; Kinchen et al., 2018; Shoshkes-Carmel et al., 2018; Thomson et al., 2018). In particular, IMSC expressing SOX6, CD142 and Wnt pathway genes appear to localize to intestinal crypts and can support epithelial stem cell function(Kinchen et al., 2018), while a population of GLI1-expressing IMSC localized in the pericrypt area has also been shown to mediate colonic stem cell renewal in a murine model (Degirmenci et al., 2018). Similarly two separate populations of fibroblast-like stromal cells (expressing *Ackr4*^+^/CD34^+^ or *Foxl1*^+^/F3^+^) in the pericrypt region have been reported to drive stem cell proliferation and gut organoid growth *in vitro*, likely via production of ligands/agonists of the Wnt and BMP pathways (Aoki et al., 2016; Shoshkes-Carmel et al., 2018; Stzepourginski et al., 2017; Thomson et al., 2018). However, it is currently unclear to what extent these different IMSC populations exert redundant functions or play different roles in gut tissue health and disease states.

The *Map3k2* gene encodes a serine/threonine protein kinase belonging to the MAP3K superfamily (Cheng et al., 2000; Guo et al., 2002; Su et al., 2001; Sun et al., 2001; Zhang et al., 2006). *Map3k2* signaling is triggered by a range of soluble factors including FGF, PDGF, TGF-β, TNF-α and IL-1β, leading to activation of downstream effectors ERK1/2, JNK, p38 and ERK5 via their respective MAPK kinases (Chang et al., 2011; Cheng et al., 2005; Greenblatt et al., 2016; Sun et al., 2001; Tsioumpekou et al., 2016; Zhang et al., 2006). *Map3k2* is constitutively expressed in multiple mouse organs including various cell types that are resident in the gut (Guo et al., 2002; Han et al., 2018).

In this report, we used scRNA-seq in combination with flow-cytometry and qRT-PCR to identify a novel population of CD81^+^ sub-crypt intestinal stromal cells that depend on a specific MAP3K2 signaling cascade to augment expression of Wnt agonist *Rspo1*. We further demonstrate that this signaling axis is critically required for rapid regeneration of damaged intestine via induction of epithelial stem cell proliferation and survival in the colon, leading to efficient host protection against DSS-induced colitis. Together, our data uncover a novel population of stromal cells that maintains an epithelial stem cell niche capable of efficient regeneration of the acutely damaged intestine.

## Results

### *Map3k2* Protects Mice against DSS-Induced Colitis

*Map3k2* is expressed in the gut epithelium, stromal and hematopoietic compartments, but germline deficiency of *Map3k2* exerts minimal effect on normal mouse growth (Chang et al., 2011; Guo et al., 2002). To investigate the role of *Map3k2* in intestinal inflammation, we used an acute colitis model in which co-housed littermates of wild type (WT) and *Map3k2^-/-^* mice were administered 2% DSS in their drinking water for 7 days followed by regular drinking water for a further two days thereafter. Within 5-6 days of DSS intake, we observed that *Map3k2^-/-^* mice suffered greater loss of body weight (**Figure 1A**) and worse diarrhea (**Figure 1B**) than did WT mice. Endoscopic examination after DSS treatment also revealed more severe bleeding, ulceration, and colonic swelling in *Map3k2^-/-^* mice than in WT mice (**Figure 1C**). Furthermore, *Map3k2^-/-^* mice also exhibited significantly shorter colon length (**Figure 1D**) as well as more severe epithelial damage and greater leukocyte infiltration than were observed in WT mice on day 9 after DSS treatment (**Figure 1E**). These data reveal that *Map3k2* plays an important role in host protection against acute intestinal tissue damage.

**Figure 1.**
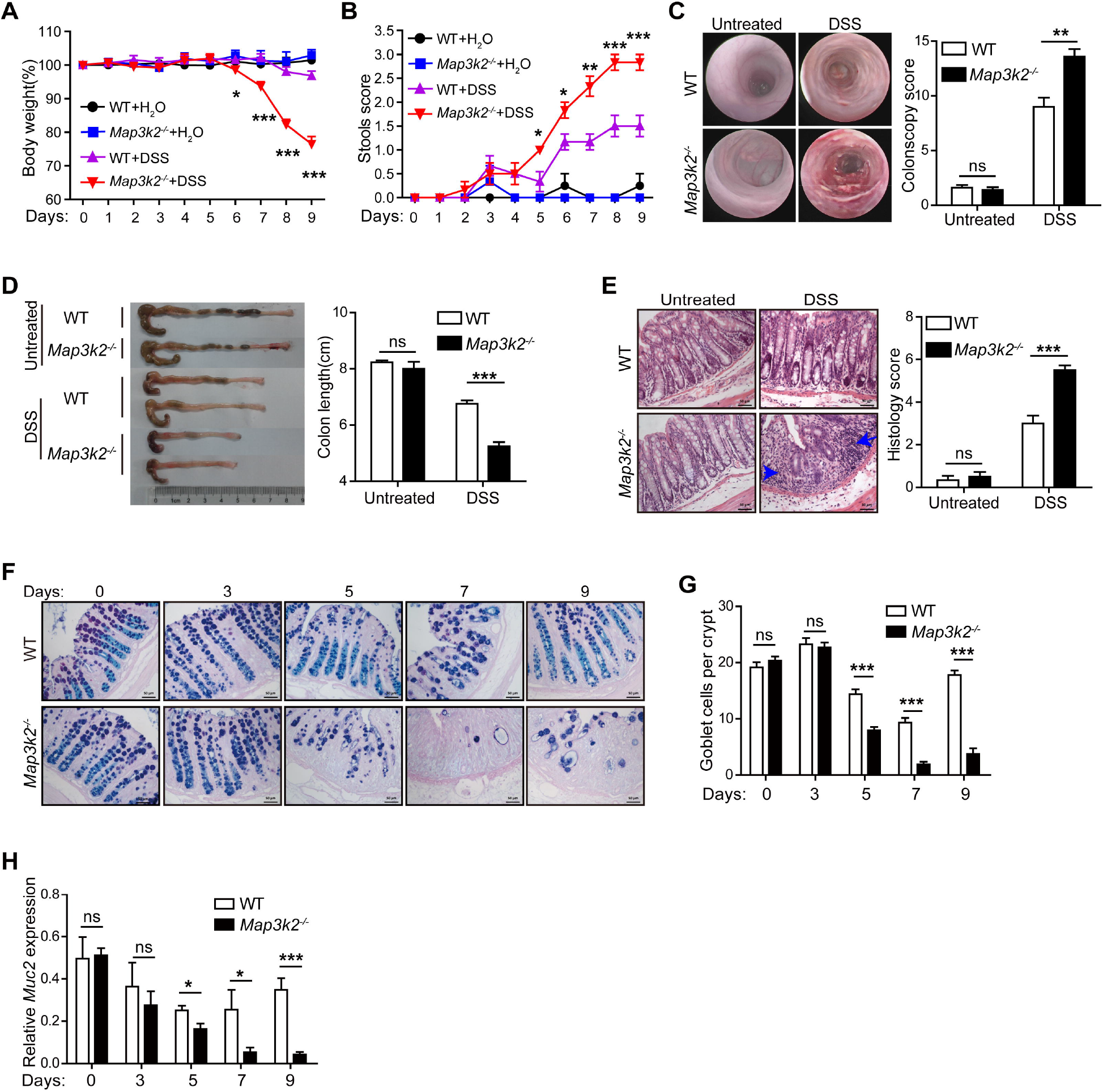
*Map3k2* Protects Mice from DSS-Induced Colitis. (A) WT mice (n=6) and *Map3k2^-/-^* co-housing littermates (n=6) were administered 2% DSS in drinking water for 7 days followed by regular drinking water for a further 2 days. In the untreated group, WT mice (n=4) and *Map3k2^-/-^* littermates (n=3) were provided with regular drinking water for entire the duration of the experiment. Body weights were recorded daily. Shown is a representative graph of three independent experiments. (B) Daily stool scores for mice treated as described in panel (A). (C) Representative colonoscopy pictures of WT and *Map3k2^-/-^* mice before and after 7 days DSS administration in drinking water (left panels) and bar graph quantification of the corresponding pathology scores (right panels) (each n=5). (D) Colon length in mice treated as described in panel (A) before sacrifice on day 9. Bar graphs show colon lengths within each experimental group (right panels). (E) Hematoxylin and eosin (H&E) staining of colon (left panels) and histological assessment of the mucosa (right panels) in mice treated as described in panel (A) before sacrifice on day 9. Arrowheads indicate infiltrating leukocytes. Pictures are representative of three independent experiments. (F) WT and *Map3k2^-/-^* co-housing littermate mice were administered 2% DSS in drinking water and then sacrificed on day 0, 3, 5, or 7 (n=3-4 mice/time-point). A separate group of mice was subjected to the same regimen for 7 days then transferred to regular drinking water for 2 days prior to sacrifice (Day 9) (n=3). Shown are representative AB/PAS staining of WT and *Map3k2^-/-^* mouse colons. (G) Goblet cell number per crypt in mice treated as described in panel (F) (20 crypts were counted in total from n=3 animals per time point). (H) mRNA expression levels of goblet cell marker gene *Muc2* in colon tissue from mice treated as described in panel (F). Data shown are normalized to *Hprt* expression level. Error bars indicate mean ± SEM (*p<0.05, **p<0.01, ***p<0.001 by unpaired Student’s t test).

### *Map3k2* is Required for Intestinal Epithelium Regeneration *In Vivo*

Given the more severe colonic damage induced by DSS administration in *Map3k2^-/-^* mice, we next assessed which specific cell types might be associated with the pathology observed in these animals. While baseline goblet cell number and capacity for mucin production were comparable in colon from both WT and *Map3k2^-/-^* mice, after 5 days DSS treatment goblet cell number were substantially reduced and spatial distribution was altered in colonic tissue from *Map3k2^-/-^* animals (**Figure 1F**). These features were even more pronounced by day 7-9, at which point the *Map3k2^-/-^* mice almost entirely lacked mucin producing cells in the gut epithelium (**Figures 1F and 1G**). Consistently, mRNA levels of the goblet cell-specific marker *Muc2* were dramatically reduced in *Map3k2*-deficient colon comparing to WT colon that were subjected to the same DSS treatment (**Figure 1H**). Ki67 staining also revealed that proliferation of colonic epithelial cells in response to DSS treatment was dramatically reduced in *Map3k2^-/-^* mice compared with WT mice (**Data not shown**). *Reg4*^+^ Deep Crypt Secreting (DCS) cells also displayed hallmarks of increased damage in *Map3k2*-deficient colon (**Data not shown**). Together, these data demonstrated that multiple lineages of intestinal epithelial cells were severely disrupted in the colon of *Map3k2^-/-^* mice, suggesting that this kinase may be critical for stem cell-mediated epithelial regeneration following acute injury.

### *Map3k2* Plays a Critical Role in Maintaining Intestinal Stem Cell Frequency

Intestinal stem cell (ISC) proliferation and differentiation are crucial for regeneration of the gut epithelium (Barker et al., 2010). Given the severity of epithelial damage observed in *Map3k2^-/-^* mice after DSS treatment, we next examined whether ISC survival and differentiation were defective in the absence of the *Map3k2* enzyme. We therefore examined colonic expression levels of the archetypal ISC genes *Lgr5* (Barker et al., 2007), *Ascl2* (Schuijers et al., 2015) and *Hopx* (Takeda et al., 2011) by qRT-PCR, which suggested a greater loss of this cell type in DSS-exposed intestinal tissue from *Map3k2*-deficient mice relative to WT animals (**Figure 2A**). To confirm these results, we next established Lgr5-EGFP reporter mice both on WT (WT-*Lgr5*-EGFP) and *Map3k2*^-/-^ (*Map3k2^-/-^-Lgr5*-EGFP) backgrounds to facilitate better comparison of the number of *Lgr5*-EGFP^+^ cells present both before and after DSS treatment. In the absence of DSS treatment, numbers of *Lgr5*-EGFP^+^ cells were comparable between colons from WT and *Map3k2^-/-^* mice, but DSS-induced damage consistently led to a greater reduction in *Lgr5*-EGFP^+^ stem cells in *Map3k2^-/-^* animals, whether assessed by flow cytometry (**Figures 2B and 2C**) or imaging quantification (**Figures 2D and 2E**). Together, these findings demonstrate that *Map3k2* protects mice against acute DSS-induced colitis by maintaining the ISC compartment.

**Figure 2.**
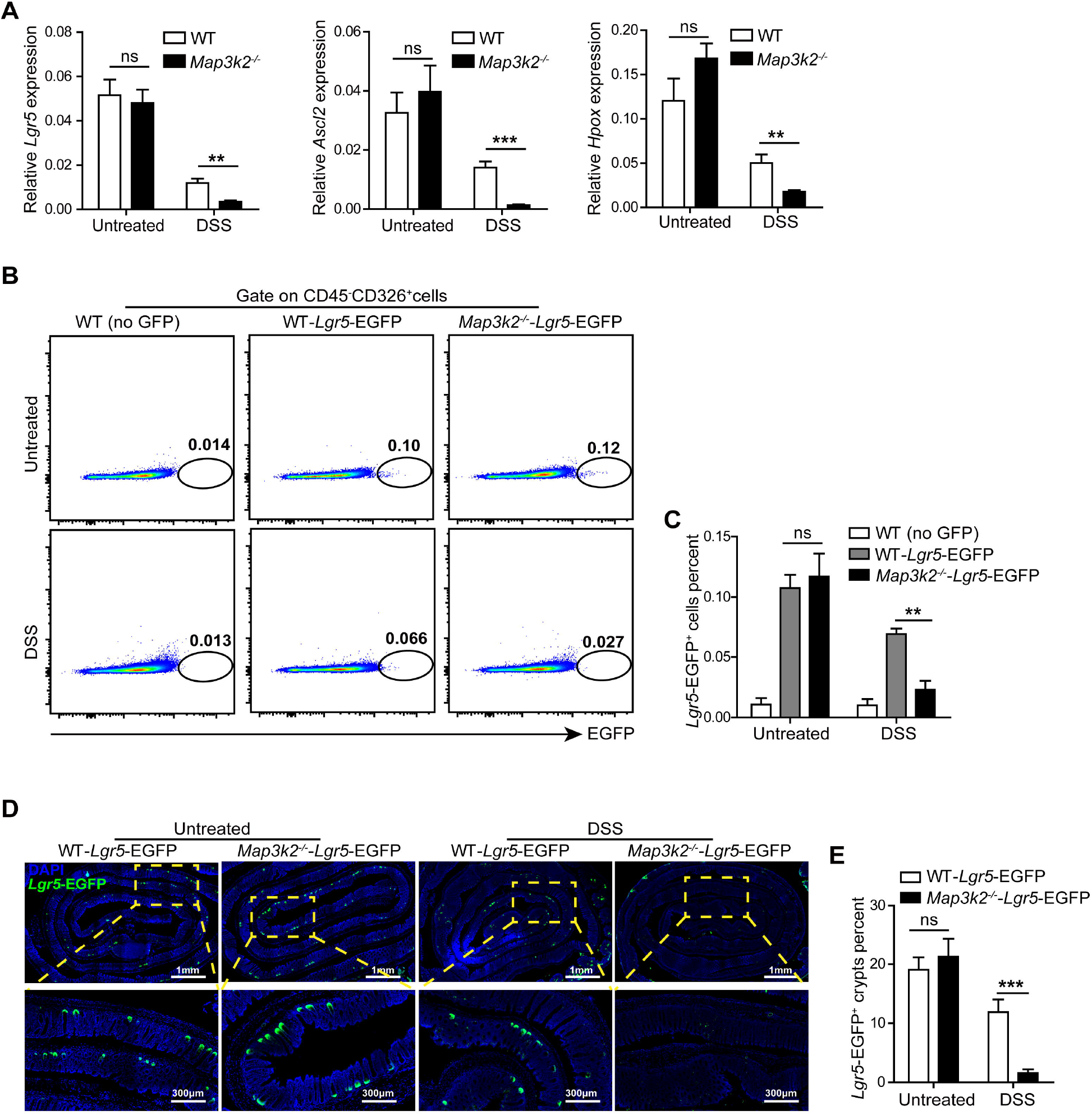
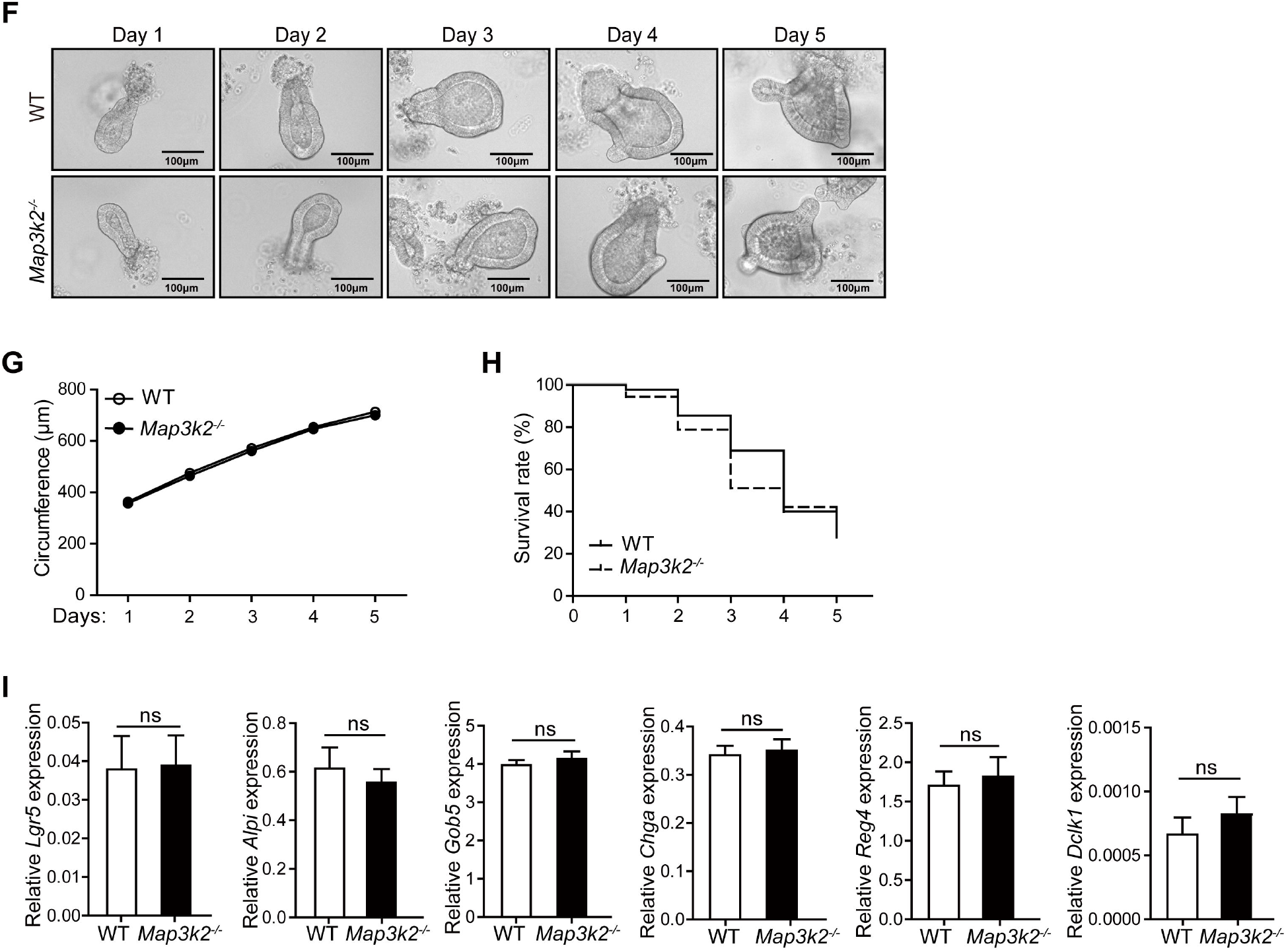
*Map3k2* Maintains Intestinal Stem Cell Frequency Following Tissue Damage. (A) *Lgr5, Ascl2* and *Hopx* mRNA expression levels in colon tissue from WT and *Map3k2^-/-^* littermate mice (each n=6) after administration of 2% DSS in drinking water for 5 days. Untreated mice (n=6) were provided with regular drinking water throughout. Data shown are normalized to *Hprt* and representative of two independent experiments. (B) Proportions of EGFP^+^ cells within the total CD45^-^CD326^+^ (Epcam) colonic epithelial cell compartment as determined by flow cytometry in WT (without GFP), *Lgr5*-EGFP, and *Map3k2^-/-^*-*Lgr5*-EGFP mice (n=4) after 5 days treatment with 2% DSS in drinking water (or regular drinking water control; n=3). (C) Quantitation of EGFP^+^ cells within the total CD45^-^CD326^+^ population of colonic epithelial cells from mice treated as described in panel (B). (D) Lgr5-EGFP and *Map3k2^-/-^-Lgr5-EGFP* mice were administered 2% DSS in drinking water for 5 days (or regular drinking water only / untreated). After 5 days, whole colons were dissected for swiss roll sectioning and stained with DAPI for visualization of GFP^+^ crypts using a Leica SP8 microscope. Dashed orange box inserts showed the representative area in each panel. (E) Quantitation of *Lgr5*-EGFP^+^ crypts in untreated and DSS-challenged mouse colons. The percentage of EGFP^+^ crypts in per 40 crypts were counted (n=8). Shown are representative pictures from two independent experiments. (F) Bright-field images of colonic crypts from WT and *Map3k2^-/-^* mice after 1-5 days culture in matrigel with Wnt3a, EGF, Noggin and R-spondin1 as indicated. Pictures shown are representative of three independent experiments. (G) Statistical analysis of average colonoid circumference (n=8) as derived from WT and *Map3k2^-/-^* mice. (H) Statistical analysis of average colonoid frequency (survival rate) (n=6) as derived from WT and *Map3k2^-/-^* mice. (I) WT and *Map3k2*-deficient colonoid mRNA expression level of *Lgr5* (intestinal stem cell marker), *Alpi* (enterocyte marker), *Gob5* (goblet cell marker), *Chga* (enteroendocrine cell marker), *Reg4* (*Reg4*^+^ deep crypt secretory [DCS] cells) and *Dclk1* (tuft cell marker) after 5 days culture in matrigel with Wnt3a, EGF, Noggin and R-spondin1 (n=6). Data shown are normalized to *Hprt* expression level. Error bars indicate mean ± SEM (**p<0.01, ***p<0.001 by unpaired Student’s t test).

### *Map3k2* is Dispensable for ISC Proliferation and Differentiation

The above data suggested that *Map3k2* might play a critical role in ISC survival, proliferation and/or differentiation into mature gut epithelial lineages including goblet cells. To investigate this possibility, we isolated intestinal crypts from WT and *Map3k2^-/-^* mice for *ex vivo* culture (Sato et al., 2009). To our surprise, there was no difference in growth rates between organoids derived from WT and *Map3k2^-/-^* animals (**Figures 2F-2H**). Furthermore, qRT-PCR analyses confirmed there was no difference in expression of hallmark genes of intestinal stem cells, enterocytes, goblet cells, enteroendocrine cells, *Reg4*^+^ DCS cells, or tuft cells (**Figure 2I**). These data suggested that *Map3k2* is not intrinsically required for ISC growth, survival and differentiation, hence the defects observed in *Map3k2^-/-^* mice *in vivo* are likely due to impairment of other cell types surrounding the stem cells and enterocytes.

### *Map3k2* Expression in Hematopoietic Cells is Not Required for Protection Against Colitis

Multiple hematopoietic and non-hematopoietic lineages are required to protect the gut against tissue damage. Since *Map3k2* is ubiquitously expressed in almost all types of intestinal cells, we postulated that the severe colonic damage observed in *Map3k2^-/-^* mice could be due to functional impairment of one or many different cell types in the gut (colonic epithelial cells, stromal cells, or hematopoietic leukocytes). We therefore performed a series of bone marrow transfers, using cells from either WT or *Map3k2^-/-^* donor mice to reconstitute lethally irradiated WT or *Map3k2^-/-^* recipient animals prior to induction of DSS colitis. Severity of inflammation was comparable in WT recipient mice irrespective of whether the bone marrow grafts were derived from WT or *Map3k2-* deficient donors, as measured by loss of body weight (**Figure S1A**), diarrhea severity score (**Figure S1B**), colon length (**Figure S1C**), or epithelial integrity (**Figure S1D**). We were also unable to detect any difference in *Lgr5* or *Gob5* expression in DSS-treated intestine from WT mice that had received grafts of either type (**Figure S1E**), suggesting that expression of *Map3k2* in hematopoietic cells was not required for protection against acute tissue damage. These findings therefore indicated that *Map3k2* expression in non-hematopoietic epithelium or tissue stroma may be critical for tissue repair after DSS-induced injury. Indeed, transfer of WT donor bone marrow into *Map3k2^-/-^* recipient mice was unable to rescue these animals from severe colitis (**Figures S1F-S1I**), and lack of *Map3k2* function only in non-hematopoietic cells was sufficient to replicate the loss of colonic epithelial stem cells and goblet cells conferred by global gene deletion (**Figure S1J**).

### *Map3k2* is Essential for Wnt Signaling in the Colon

To better understand the molecular basis of *Map3k2*-mediated protection against DSS-induced colitis, we next isolated and sequenced total RNA from the colons of WT and *Map3k2^-/-^* mice after one day exposure to DSS or normal drinking water. In order to identify key *Map3k2* target genes and associated signaling pathways, genes that were differentially expressed between the paired samples were determined using Gene Set Enrichment Analysis (GSEA). Using this approach, we detected significant up-regulation of chemokine genes in *Map3k2*-deficient colon after DSS exposure, consistent with the increased inflammatory infiltrate observed in these animals, whereas genes involved in MAPK signaling were down-regulated as expected (**Data not shown**). However, when we assessed expression levels of genes associated with intestinal stem cell survival and maintenance, we observed that *Map3k2^-/-^* mice exhibit potent down-regulation of Wnt signaling genes (**Figure 3A**) but not the Notch or EGF pathways (**Figure 3B**). Furthermore, while expression of Wnt pathway genes in untreated colon tissue was comparable between WT and *Map3k2^-/-^* animals, DSS exposure was associated with a dramatic down-regulation of key Wnt pathway-associated genes such as *Axin2, Fzd8, Nr4a3* and *Rspo1* in *Map3k2*-deficient colon (**Figures 3A and 3C**). Furthermore, the overall levels of the ligands, receptors, and mediators of the Wnt pathway were all down regulated in *Map3k2*-deficient colon after DSS treatment (**Figure 3D**), strongly indicating that *Map3k2* critically regulates Wnt signaling in order to protect the colon against DSS-induced epithelial damage.

**Figure 3.**
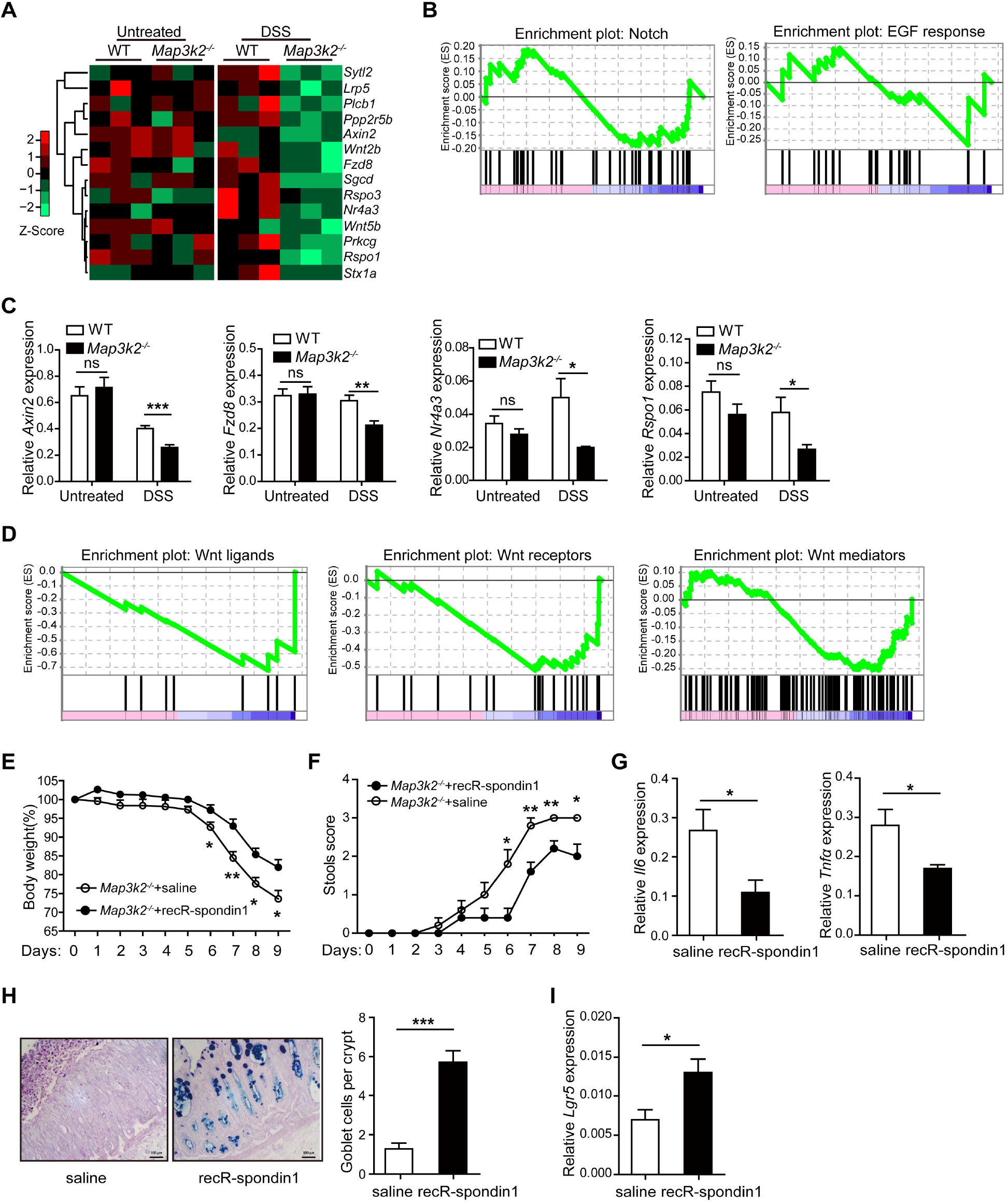
Wnt agonist R-spondin1 Rescues *Map3k2^-/-^* Mice from DSS-Induced Severe Colitis. (A) WT and *Map3k2^-/-^* mice (each n=3) were administered 2% DSS or regular drinking water (untreated) for one day before total colon tissue RNA was isolated for RNA-seq. Heat-map of the RNA-seq data showing Wnt pathway genes that were differentially expressed between WT and *Map3k2*-deficient mouse colons. (B) Gene Set Enrichment Analysis (GSEA) of the Notch pathway and EGFR response target genes in WT and *Map3k2*-deficient colons was performed and is presented as an enrichment plot. (C) qRT-PCR analysis of *Axin2, Fzd8, Nr4a3* and *Rspo1* mRNA levels in colonic tissue from mice treated as described in panel (A). Data are normalized to *Hprt* expression level. (D) Gene Set Enrichment Analysis of Wnt ligands, Wnt receptors, and Wnt signal mediators in WT and *Map3k2* deficient colons from mice treated as described in panel (A). (E) *Map3k2^-/-^* mice (n=5/group) were administered 2% DSS in drinking water for 7 days followed by regular drinking water for 2 days. Human recombinant R-spondin1 was injected at a daily dose of 5μg/mouse (from day 1-9). Control mice received saline injection only. Body weight was monitored daily. (F) Daily stool scores from mice treated as described in panel (E). (G) *Il6* and *Tnfa* mRNA expression levels in colon tissue from mice treated as described in panel (E) before sacrifice on day 9. Data shown are normalized to *Hprt* expression level. (H) Representative Alcian Blue/Periodic Acid Schiff (AB/PAS) staining of colons from mice treated as described in panel (E) before sacrifice on day 9 (left panels). Goblet cell number per crypt is shown on the right (n=5 mice/group, 30 crypts counted per group). (I) *Lgr5* mRNA expression in colon tissue from mice treated as described in panel (E) before sacrifice on day 9. Data shown are normalized to *Hprt* expression level. Error bars indicate mean ± SEM (*p<0.05, **p<0.01, ***p<0.001 by unpaired Student’s t test).

Among the Wnt-related genes identified above, *Rspo1* was the most significantly downregulated in *Map3k2*-deficient colon (**Figures 3A and 3C**), consistent with the reports that *Rspo1* plays an essential role in maintaining the ISC niche (Kim et al., 2005; Ootani et al., 2009; Sato et al., 2009; Zhao et al., 2007; Zhou et al., 2013). Accordingly, we observed that reducing the concentrations of R-spondin1 in the *in vitro* culture media led to dose-dependent decreases in growth and survival of intestinal organoids (**Figures S2A-S2C**), strongly suggesting that peri-crypt concentrations of R-spondin1 may be equally critical for proliferation and survival of gut epithelial stem cells *in vivo*.

### Recombinant R-spondin1 Rescues *Map3k2^-/-^* Mice from DSS-Induced Severe Colitis

To determine whether *Map3k2* is crucial for R-spondin1 induction and protection against DSS-induced intestinal damage, we next performed a rescue experiment using human recombinant (rec) R-spondin1. Specifically, *Map3k2^-/-^* mice were administered 2% DSS in drinking water with daily intraperitoneal injection of 5 μg recR-spondin1 or saline-only control. As shown in **Figures 3E and 3F**, recR-spondin1 markedly reduced colitis severity in *Map3k2^-/-^* mice as indicated by less pronounced weight loss (**Figure 3E**) and milder symptoms of diarrhea (**Figure 3F**). Expression of inflammatory cytokines such as *Il6* and *Tnfα* was also dramatically reduced in recR-spondin1-treated mice colon tissues (**Figure 3G**). We further observed a partial rescue of goblet cell numbers as assessed by AB/PAS staining (**Figure 3H**), together with increased mRNA expression of *Lgr5* (**Figure 3I**). Together, these data strongly suggest that the protective role of *Map3k2* in DSS-induced colitis is mediated via up-regulation of *Rspo1*.

### *Map3k2* Regulates Intestinal Mesenchymal Stromal Cell Induction of R-spondin1

R-spondin1 is crucial for optimal activation of the Wnt pathway, which plays well-recognized roles in maintaining gut integrity and repairing acute tissue damage (Binnerts et al., 2007; de Lau et al., 2014; Kim et al., 2005; Kretzschmar and Clevers, 2017; Pinto et al., 2003; Yan et al., 2017). However, the cellular sources of Wnt agonist R-spondin1 in the damaged gut have not been clearly defined. In order to uncover the key cellular source of *Map3k2*-dependent *Rspo1* expression in the DSS-damaged gut, we used flow cytometry to sort CD326^+^CD45^-^ intestinal epithelial cells (IECs), CD45^+^CD326^-^ hematopoietic cells (CD45^+^), and CD326^-^CD45^-^CD31^-^gp38^+^ intestinal mesenchymal stromal cells (gp38^+^ stroma) from untreated and DSS-challenged mouse colon (**Figure 4A**). In addition, we used anti-CD90 antibody to sort the gp38^+^ stromal fraction into CD90^+^ IMSC and CD90^-^ IMSC subsets before assessing *Rspo1* expression by qRT-PCR. As predicted, we were unable to detect *Rspo1* expression in sorted IEC, whereas CD45^+^ hematopoietic cells and CD90^-^ IMSC expressed only low levels of *Rspo1* (**Figure 4B**). In contrast, CD90^+^ IMSC displayed marked production of *Rspo1* and higher expression of *Map3k2* than did any other colonic cell population analyzed (**Figures 4B and 4C**). Importantly, we further observed that *Rspo1* expression in CD90^+^ IMSC was significantly up-regulated following DSS treatment (**Figure 4D**), but only in the presence of functional *Map3k2* (**Figure 4D**). These data demonstrate that CD90^+^ IMSC are likely the key stromal population responsible for *Map3k2*-mediated gut tissue repair.

**Figure 4.**
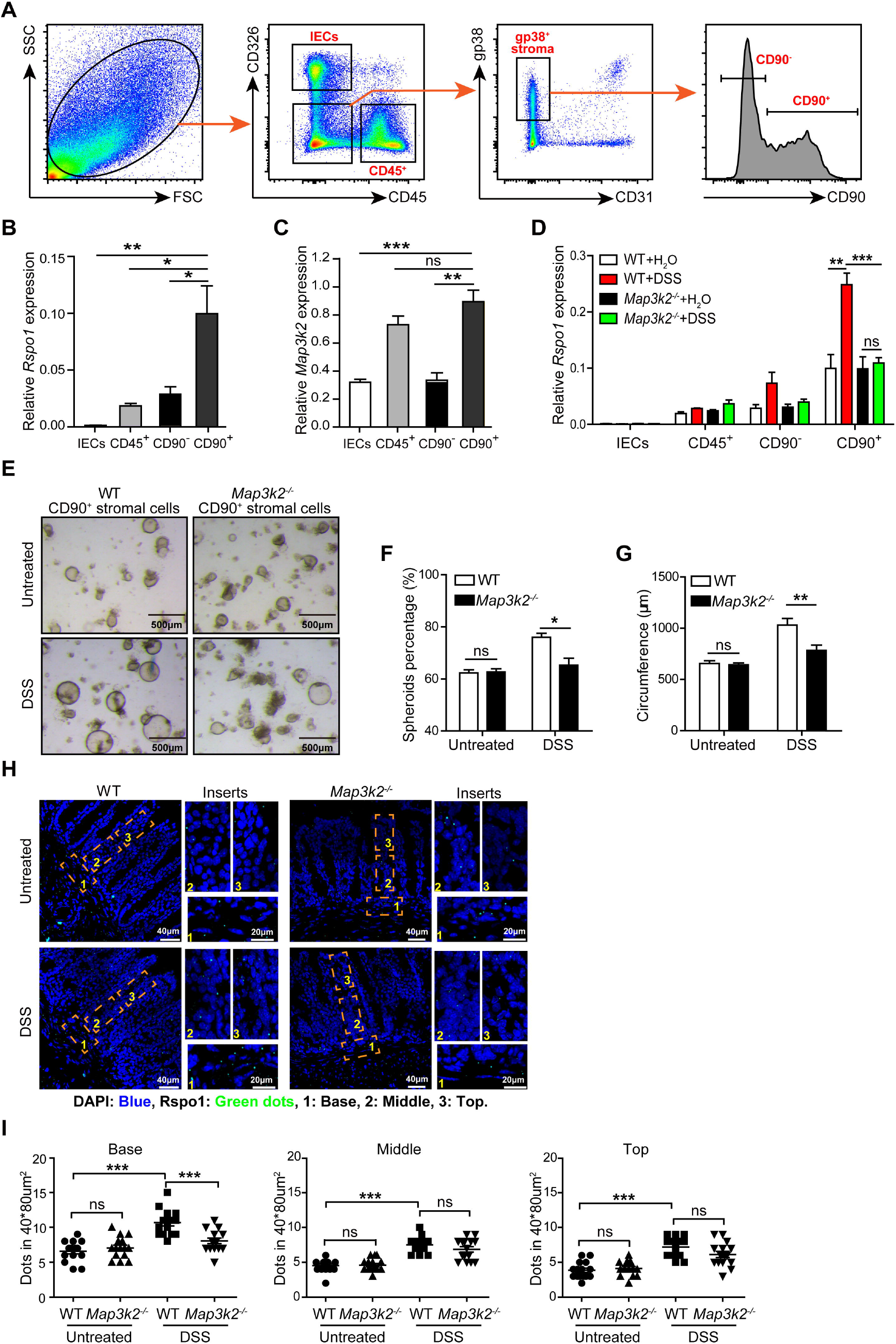
*Map3k2*-Deficient CD90^+^ Stromal Cells Display Impaired *Rspo1* Induction. (A) Representative flow-cytometry plots showing the percentages of CD326^+^ intestinal epithelial cells (IEC), CD45^+^ leukocytes, CD90^-^ intestinal mesenchymal stromal cells (CD45^-^CD31^-^CD326^-^gp38^+^CD90^-^), and CD90^+^ intestinal mesenchymal stromal cells (CD45^-^CD31^-^CD326^-^gp38^+^CD90^+^) present in WT colon. Shown are representative plots from three independent experiments. (B) *Rspo1* mRNA expression in the indicated cell types sorted as described in panel (A). Data shown are normalized to *Hprt* expression level (n=4). (C) *Map3k2* mRNA expression in the indicated cell types sorted as described in panel (A). Data shown are normalized to *Hprt* expression level (n=4). (D) Relative *Rspo1* mRNA expression in CD326^+^ intestinal epithelial cells (IEC), CD45^+^ leukocytes, CD90^-^ intestinal mesenchymal stromal cells (CD45^-^CD31^-^CD326^-^gp38^+^CD90^-^), and CD90^+^ intestinal mesenchymal stromal cells (CD45^-^CD31^-^CD326^-^gp38^+^CD90^-^) FACS-sorted from WT and *Map3k2* deficient colons (n=4/group) after 1 day treatment with 2% DSS (or water only/untreated control). Data shown are normalized to *Hprt* expression level and are representative of two independent experiments. (E) Bright-field images of matrigel-embedded colonic crypts cultured together with CD90^+^ intestinal mesenchymal stromal cells in the presence of EGF, Noggin and Wnt3a for 3 days. CD90^+^ intestinal mesenchymal stromal cells were sorted by flow cytometry from WT and *Map3k2^-/-^* mouse colons after 1-day DSS treatment (or water only/untreated control). (F) Spheroids percentages among total colonoids derived after 3 days culture under the conditions described in panel (E) (n=3/group). (G) Statistical analysis of average colonoid circumference after 3 days culture under the conditions described in panel (E). (H) *Rspo1* mRNA detection using fluorescence *in situ* hybridization (FISH)-based RNAScope analysis of colons from WT and *Map3k2^-/-^* mice after 1-day exposure to 2% DSS (or normal drinking water control). Dashed orange box inserts (40μm*80μm) in each panel indicate the base (1), middle (2), and top (3) sections of the sub-epithelial area along the crypt axis. (I) Quantification of *Rspo1* mRNA expression by FISH (RNAScope) as described in panel (G). Fluorescence signals were counted within defined box areas covering 15 individual sub-epithelial regions along the crypt axis from base to top. Shown are summary data of *Rspo1* mRNA fluorescence signal per 40*80μm^2^ area of the crypt axis for the base, middle and top regions. Error bars indicate mean ± SEM (*p<0.05, **p<0.01, ***p<0.001 by unpaired Student’s t test).

In order to confirm that CD90^+^ IMSC are physiologically important for supporting the growth of intestinal epithelial stem cells, we performed a stroma-organoid co-culture experiment using isolated CD90^+^ IMSC or CD90^-^ IMSC co-cultured together with colonic crypts in the absence of exogenous R-spondin1. Organoids displayed only limited growth when cultured in isolation, whereas co-culture with CD90^+^ IMSC led to a marked increase in overall size and the percentages of spheroids (**Figures S2D and S2E**). Intriguingly, CD90^-^ IMSC not only failed to promote growth but actually appeared to inhibit organoid expansion when compared with cultures that lacked any stroma (**Figures S2D and S2E**). Having already established that *Map3k2*-deficient CD90^+^ IMSC was poor inducers of R-spondin1, which was essential for growth and survival of intestinal organoids (**Figure S2A**), we next assessed whether CD90^+^ IMSC were still able to support organoid growth after deletion of the *Map3k2* gene. Using the same stroma-organoid co-culture approach, we observed that CD90^+^ IMSC from WT mouse colon were potent inducers of colonic organoid growth after DSS exposure, forming more spheroids with greater circumference than could be established by their *Map3k2*-deficient counterparts (**Figures 4E-4G**). In contrast, in animals that did not receive DSS, colonic CD90^+^ IMSC derived from either WT or *Map3k2^-/-^* mice displayed similar capacities to support organoid growth (**Figures 4E-4G**). All together, these results strongly suggest that the protective function of *Map3k2* in the gut is mediated by CD90^+^ but not CD90^-^ IMSC.

### *Map3k2*-Regulated CD90^+^ IMSC Are Localized to the Colonic Sub-Crypt

To determine the anatomical location of *Map3k2*-regulated CD90^+^ IMSC in the gut, we next performed immunofluorescent staining using stromal specific markers together with an anti-CD90 antibody. As shown in **Figure S3A**, we observed characteristic CD90 staining of the lamina propria T cells or innate lymphoid cells (ILCs), but also detected a specific population of CD90^+^gp38^+^ cells located just below the crypts where LGR5-EGFP^+^ stem cells were also situated as described in a recent study (**Figure S3A**) (Karpus et al., 2019). However, despite lacking the epithelial marker CD326 and leukocyte antigen CD45, further analysis of this population revealed a monolayer-like structure that is not typical of mesenchymal stroma, and subsequent whole-mount anti-CD90 staining identified this as a nexus of lymphatic vessels that co-stained for the endothelial marker LYVE1 (**Figures S3B and S3C**, Video 1 and Video 2). We therefore reverted to flow cytometry in order to perform a systemic analysis of which colonic subsets express CD90. IEC and gp38^-^ stromal cells were found to express little CD90, and only a very small fraction of CD45^+^ cells expressed this molecule (likely the T-cells and ILCs; **Figures S3D and S3E**). Between 30-50% of blood endothelial cells (BEC) and gp38^+^ IMSC were found to express moderate levels of CD90, whereas high-level expression of this marker was restricted to lymphatic endothelial cells (LEC) (**Figure S3E**). Indeed, when assessed by flow cytometry, LEC expression levels of CD90 were approximately 10-fold higher than those detected for *Map3k2*-regulated gp38^+^ IMSC (**Figure S3E**). While we anticipate that CD90^+^ IMSC are situated in a similar location to LEC *in vivo*, the levels of CD90 expressed by lymphatic endothelial cells prevented *Map3k2*-dependent CD90^+^ IMSC from being confidently located by microscopy. Consequently, we hereafter refer to CD90^+^ IMSC as ‘CD90 medium-high’ (CD90^mid^) cells.

Although CD90 staining did not reveal the identity of the *Map3k2*-regulated IMSC, a sub-crypt location for these cells could be predicted due to the profound effect of *Map3k2* deletion on the ISC niche after DSS challenge. To examine this possibility, we used the fluorescence *in situ* hybridization (FISH)-based technique RNAScope to detect *Rspo1* mRNA in tissue sections, which revealed that DSS exposure in drinking water was able to induce *Rspo1* expression in the colonic epithelium of WT mice (**Figure 4H**). Quantitation of *Rspo1* expression levels showed marked induction in the base areas of the sub-epithelium after DSS-treatment, but this was blunted in colon from *Map3k2^-/-^* mice (**Figure 4I**). In contrast, *Rspo1* expression in the middle and top layers of the sub-epithelial region was comparable in colons from both WT and *Map3k2*-deficient animals, irrespective of DSS treatment (**Figure 4I**). Together, these data suggest that *Map3k2*-mediated *Rspo1* expression defines a novel ISC niche located at the bottom of the crypts.

### Single-cell RNA-Seq Reveals that CD90^mid^CD34^+^CD81^+^ IMSC Require *Map3k2* for DSS-induced *Rspo1* Expression

Having determined that CD90^mid^ IMSC are likely the critical subset of *Map3k2*-regulated cells expressing *Rspo1* in response to acute gut injury, we next sought to uncover the precise features of this stromal population using droplet-based scRNA-seq. First, we sorted CD90^mid^ IMSC from DSS-treated colonic lamina propria by gating out hematopoietic cells (CD45^+^), epithelial cells (CD326^+^), endothelial cells (CD31^+^), and gp38^-^ stromal cells (gp38^-^) (**Figure S4A and S4B**), then sequenced 11,116 individual CD90^mid^ IMSC using the 10x Genomics platform. Based on unsupervised clustering using Seurat software (Butler et al., 2018), CD90^mid^ IMSC were grouped into 10 distinct subsets with specific gene expression profiles (Amir el et al., 2013) (**Figure 5A**), each of which lacked expression of genes excluded by our cell sorting strategy (CD31, CD45, CD326) (**Figure 5B**). Preliminary analysis confirmed that CD90^mid^ cells were positive for *Pdpn* (gp38), *Vim, Col1a2*, and *Ly6a*, but did not express the enteric neuronal cell marker gene *Ret* (**Figure 5B**). According to the schemes utilized in other studies (Powell et al., 2011), these 10 IMSC clusters could be tentatively assigned as myofibroblasts (cluster 4) (Mifflin et al., 2011), MHC II^+^ stromal cells (cluster 6) (Messina et al., 2017), telocytes (cluster 7) (Shoshkes-Carmel et al., 2018), mesothelial cells (cluster 8) (Rinkevich et al., 2012), interstitial cells of cajal (cluster 9) (Kinchen et al., 2018), contaminating epithelial cells (cluster 10) (Haber et al., 2017), and at least four subgroups of mesenchymal stromal cells for which no prior identifying markers have been reported (clusters 1, 2, 3, & 5) (*Figure 5A*). Among these populations, clusters 2, 5, and 8 were found to express *Rspo1* at high levels, identifying these as candidate *Map3k2*-regulated stromal cells potentially capable of responding to DSS-induced gut tissue damage (*Figure 5C*). Of these subsets, cluster 8 mesothelial cells are situated away from the crypt region and are thus unlikely to play a direct role in supporting epithelial reconstitution. We therefore focused subsequent analyses on the remaining Rspo1-expressing cell clusters 2 and 5.

**Figure 5.**
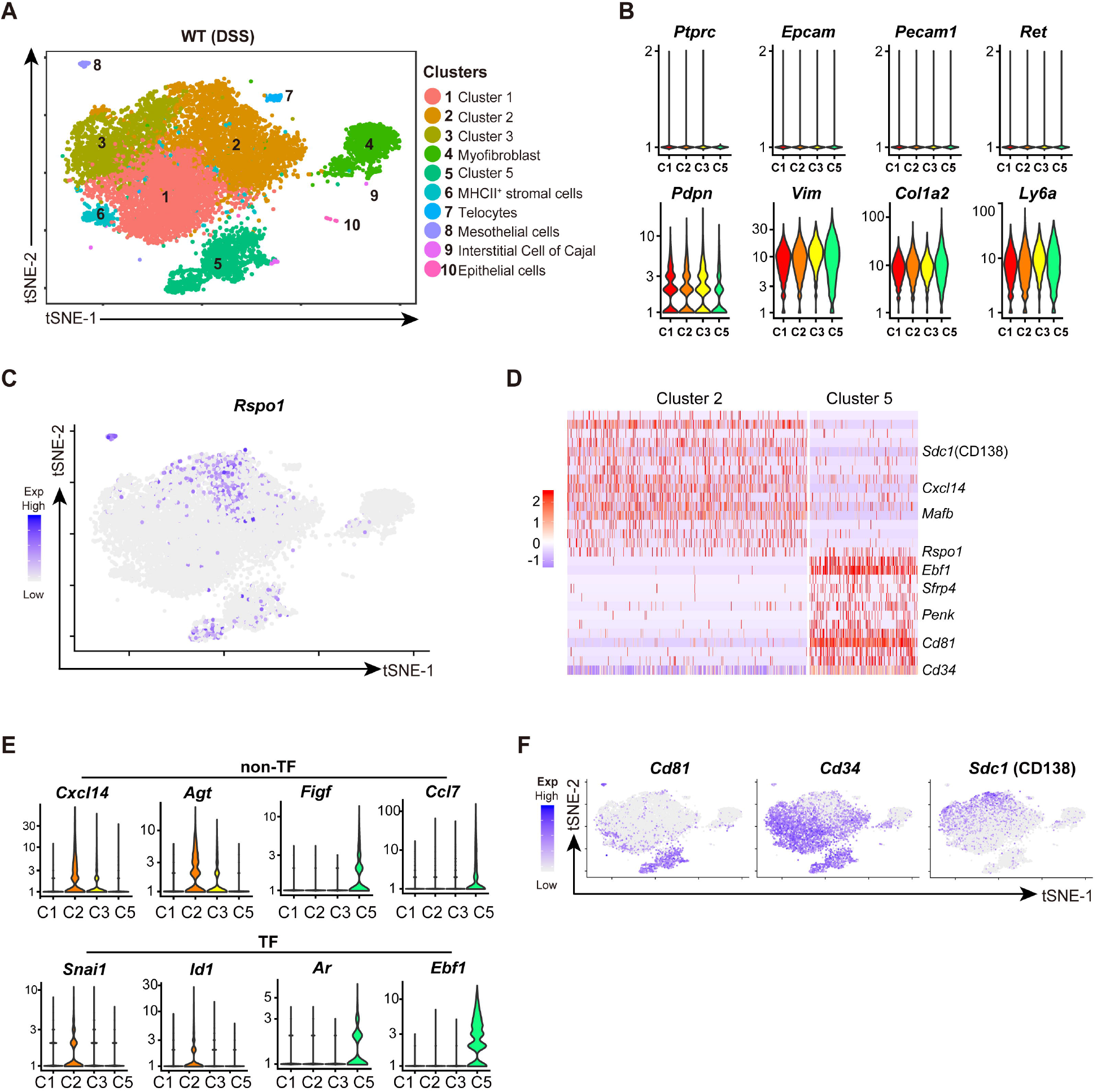
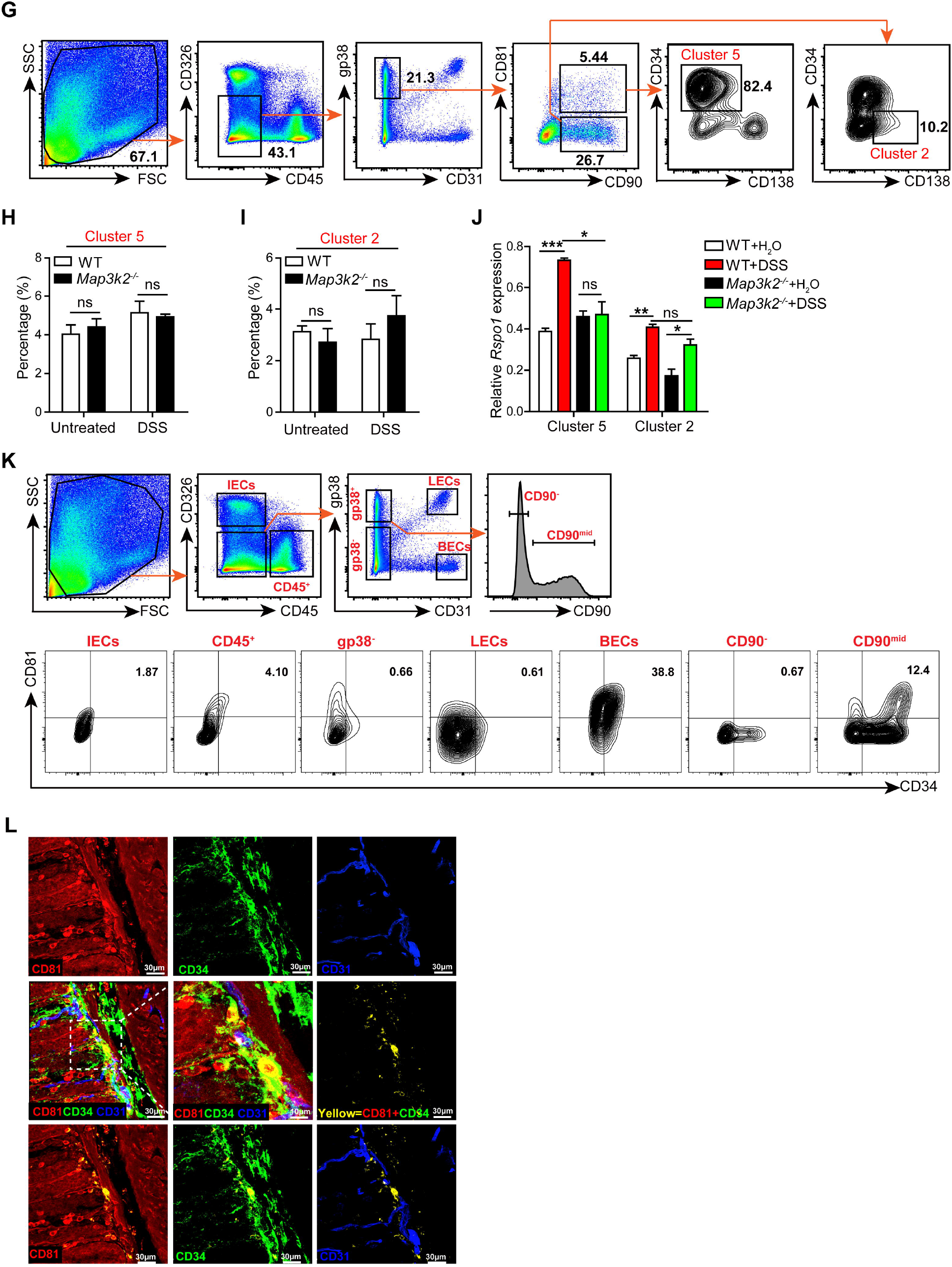
*Map3k2*-regulated Intestinal Stromal Cells are Defined by a CD90^mid^CD81^+^ Phenotype. (A) t-Distributed Stochastic Neighbor Embedding (t-SNE) plot of the scRNA-seq dataset generated by CD45^-^CD326^-^CD31^-^gp38^+^CD90^mid^ stromal cells isolated from WT mouse colon after 3 days exposure to 2% DSS in drinking water. Each dot represents an individual cell, color-coded according to cluster annotation. (B) Violin plots of the indicated genes expressed in clusters 1, 2, 3, and 5. (C) t-SNE plot showing *Rspo1* gene expression in the scRNA-seq dataset at single cell level. (D) Heat-map of genes that were significantly differentially expressed between cluster 2 and cluster 5, shown at single cell level. (E) Violin plots of highly-ranked marker genes (non-TF) and transcriptional regulators (TF) expressed in cluster 2 and cluster 5, as compared with other clusters, as indicated. (F) t-SNE plots showing *Cd81, Cd34*, and *Sdc1* gene expression in the scRNA-seq dataset at single cell resolution. (G) Representative flow cytometry plots showing percentage of cluster 5 stromal cells (CD45^-^, CD31^-^, CD326^-^, gp38^+^, C90^mid^, CD81^+^, CD34^+^, CD138^-^) and cluster 2 stromal cells (CD45^-^, CD31^-^, CD326^-^, gp38^+^, CD90^mid^, CD81^-^, CD34^-^, CD138^+^) in WT colonic lamina propria. (H) Bar graph quantitation of the percentage of cluster 5 stromal cells (CD45^-^, CD31^-^, CD326^-^, gp38^+^, CD90^mid^, CD81^+^, CD34^+^, CD138^-^) in gp38^+^ stromal cells as assessed by flow cytometry analysis in WT and *Map3k2^-/-^* mouse colons (each n=3) after treatment or not with DSS as indicated. Shown are representative plots from three independent experiments. (I) Bar graph quantitation of the percentage of cluster 2 stromal cells (CD45^-^, CD31^-^, CD326^-^, gp38^+^, CD90^mid^, CD81^+^, CD34^+^, CD138^-^) in gp38^+^ stromal cells as determined by flow cytometry analysis in WT and *Map3k2^-/-^* mouse colons (each n=3) after 2 days exposure to DSS (or water only control) as indicated. (J) Intestinal stromal cell clusters 2 and 5 were FACS-sorted from the colonic tissue of WT and *Map3k2^-/-^* mice (each n=3) after 2 days exposure to DSS (or water only control) and total RNA was extracted for detection of *Rspo1* mRNA. Data are normalized to *Hprt* expression level. (K) Flow cytometry analysis of CD34 and CD81 expression in WT colonic lamina propria showing IEC (CD326^+^), CD45^+^ leukocytes, LECs (CD45^-^CD326^-^CD31^+^gp38^+^), BECs (CD45^-^CD326^-^ CD31^+^gp38^-^), gp38^-^ cells (CD45^-^CD326^-^CD31^-^gp38^-^), CD90^-^ stromal cells (CD45^-^CD326^-^CD31^-^ gp38^+^CD90^-^) and CD90^mid^ stromal cells (CD45^-^CD326^-^CD31^-^gp38^+^CD90^mid^). Plots are representative of three independent experiments. (L) WT mouse colon was harvested and processed into swiss roll sections for staining with anti-CD81 (red), anti-CD34 (green), anti-CD31 (blue) and then visualized using a Leica SP8 microscope with 63x oil objective lens. The top row panels show the images of three separate channels as indicated. The left middle panel show an image of merged channels, and the center panel in the middle row shows a magnified image of the area from the left panel highlighted by dotted lines. The right panel in the middle row shows the CD81 and CD34 double positive staining marked with a pseudo-yellow color. The bottom row panels show CD81 and CD34 double positive cells (yellow) superimposed on each of the other channels as indicated. Pictures displayed are representative of two independent experiments. Error bars indicate mean ± SEM (*p<0.05, **p<0.01, ***p<0.001 by unpaired Student’s t test).

To investigate the properties of these two stromal cell populations, we first examined the characteristic marker genes that differentiated these into clusters 2 and 5. Differentially expressed gene analysis indicated that cluster 2 was more pro-inflammatory, as indicated by preferential expression of leukocyte/endothelial cell chemokine *Cxcl14* and the activation-linked transcription factor *Mafb*. In contrast, cluster 5 cells displayed a profile consistent with involvement in tissue remodeling, and preferentially expressed the neuro-regulator *Penk*, Wnt modulator *Sfrp4*, and cell development regulator tetraspanin *Cd81* (**Figure 5D**). Since both cluster 2 and 5 cell subsets lacked genes such as *Sox6* that were characteristic of other niche-associated stromal cells such as telocytes (Shoshkes-Carmel et al., 2018), these data suggest that the populations detected by our analyses represent two previously unknown stromal cell lineages with potential roles in regulating the intestinal stem cell niche (**Figure 5D**).

Using violin plots to compare specific gene expression within the different cell populations, we found that cluster 2 population expressed mediators typically associated with angiogenesis (*Cxcl14* and Agt), whereas cluster 5 cells were found to express genes potentially related to tissue damage and repair (*Figf* and *Ccl7)* (**Figure 5E**). Indeed, cluster 2 cells were further defined by unique expression of the transcription factors *Snai1* and Id1, while cluster 5 cells instead expressed *Ar* and *Ebf1* (**Figure 5E**), indicating that these two stromal lineages exert different functions in the colon. In order to facilitate further analysis of these populations by flow cytometry, we next examined their expression profiles of cell-surface markers, which revealed that cluster 5 cells displayed high level of CD81 and CD34 but not CD138 (*Sdc1*), whereas cluster 2 cells lacked CD81 and CD34 but were CD138 positive (**Figure 5F**). To determine which of these populations was the major *Map3k2*-regulated subset involved in gut tissue repair, we next stained colonic lamina propria single cell suspensions with a panel of lineage-defining antibodies (CD326, CD45, CD31, gp38, CD90, CD81, CD34, CD138). The results revealed that about 82.4% of CD81^+^CD90^mid^ stromal cells, which were fewer than ~4% of the total gp38^+^ stromal cells, were likely cluster 5 cells, whereas 10.2% of CD81^-^CD90^mid^ stromal cells belonged to cluster 2 (**Figure 5G**). When we sorted these two populations of *Rspo1*^+^ IMSC from unchallenged mice, we observed comparably-sized populations in both WT and *Map3k2*-deficient colon (**Figures 5H and 5I**). After DSS treatment, we detected a very small increase of cluster 5 cell frequency (CD90^mid^ CD34^+^CD81^+^ CD138^-^) in both WT and *Map3k2^-/-^* mice (**Figure 5H**), whereas cluster 2 cells (CD90^mid^CD81^-^CD34^-^CD138^+^) appeared unaffected by DSS damage (**Figure 5I**). Furthermore, basal expression of *Rspo1* gene was similar in cluster 5 cells from both WT and *Map3k2^-/-^* mice, but after DSS administration expression of *Rspo1* in this cluster was substantially upregulated only in *Map3k2*-competent cells (**Figure 5J**). In contrast, cluster 2 cells expressed comparable low levels of *Rspo1* in both WT and *Map3k2* animals, and the modest increase in *Rspo1* observed after DSS challenge appeared unaffected by the absence of functional *Map3k2* (**Figure 5J**). Together, these data demonstrate that the use of common lineage markers together with CD81 and CD34 has identified cluster 2 as the *Map3k2*-regulated stromal cells implicated in ISC niche maintenance.

If *Map3k2*-dependent CD81^+^CD34^+^ stromal cells were a genuine niche-forming population, we would expect these cells to be situated in close proximity to intestinal stem cells. Before testing this hypothesis using imaging approaches, we first validated our staining strategy by using flow cytometry to determine which colonic lamina propria populations express both CD81 and CD34. As shown in **Figure 5K**, IECs displayed only low-level expression of CD81 while completely lacking CD34, and the few CD45^+^ leukocytes that stained positive for CD81 were mostly negative for CD34. LECs were consistently negative for both markers. Around 38.8% of BECs expressed modest levels of CD34 in conjunction with CD81, but these cells could be clearly distinguished from stromal cells based on the additional expression of CD31. Finally, dual expression of CD81 and CD34 was negligible among gp38^-^ stromal cells and totally absent within the CD90^-^ stromal compartment. Taken together, these data confirmed that ~12% of the CD90^mid^ stromal population could be uniquely defined as CD81^+^CD34^+^ ‘double-positive’ cells, and could therefore be identified by image analysis for assessment of their tissue distribution within the gut.

We next used antibodies against CD81, CD34 and CD31 to perform immunofluorescence analysis of colon sections in order to determine whether CD81^+^CD34^+^ stromal cells could be identified within the sub-crypt region. While anti-CD81 antibody labelled most cells in colon (red), no staining was observed within the sub-crypt region which was instead dominated by CD34 (green) co-expression with CD31 (blue) indicative of endothelial cells (BEC, LECs) (**Figure 5L**). However, we were also able to detect CD31^-^ cells that co-expressed CD81 and CD34 (yellow) near the bottom of the crypts, where they appeared to be tightly associated with lymphatic vessels (**Figure 5L, Video 3**). These results may explain why we were previously unable to visualize these cells using only anti-CD90 staining, since this marker is expressed at 10-fold higher levels by adjacent LECs (**Figure S3E**). Taken together, these data indicate that a novel lineage of colonic mesenchymal stromal cells defined by co-expression of CD81 and CD34 (but not CD31) is situated close to the lymphatic vessels and depends on *Map3k2* to induce *Rspo1* expression following DSS-induced tissue damage. Hereafter, these cells will be referred to as *Map3k2*-regulated intestinal stromal cells (MRISC).

### Conditional Ablation of *Map3k2* in Mice Confirms its Intestinal Stromal Cell Specific Roles

To conclusively demonstrate the stromal cell specific role of *Map3k2 in vivo* for its protection of damaged colon, we next generated a *Map3k2* floxed (*Map3k2^fl/fl^*) mouse line by inserting two loxp sequences flanking the exon 5 of *Map3k2* gene (**Figure S5A**). The tissue specific deletion of *Map3k2* gene in these mice was confirmed by first crossing these mice with a T cell specific *CD4-Cre* line (Sawada et al., 1994). CD4^+^ T cells and CD19^+^ B cells were FACS sorted from control *Map3k2^fl/fl^* or *CD4-Cre:Map3k2^fl/fl^* mice for PCR genotyping, and mouse tail DNAs were extracted from WT mice as well as *Map3k2^fl/fl^* and *Map3k2^fl/+^* as controls for the PCR genotyping (**Figure S5B**). Cell lysates from CD4^+^ T cells and CD19^+^ B cells were further used for immunoblotting to verify the tissue specific *Map3k2* deletion in CD4^+^ T cells but not B cells (**Figure S5C**). These results demonstrate that the *Map3k2^fl/fl^* mice can be used for tissue specific deletion of the *Map3k2* gene.

We next bred the *Map3k2^fl/fl^* mice with a commercially available stromal cell specific transgenic line *Pdgfra-Cre^ERT^* to study the *in vivo* role of *Map3k2^fl/fl^* in the gut (Kang et al., 2010). However, we found out that this particular transgenic *Cre* line could not delete the target genes from the gut stromal cells effectively although it was reported to delete genes in stromal cells in other organs/tissues (**Data not shown**). Therefore, we generated a new stromal specific *Col1a2-Cre^ERT2^* line, in which a *Cre^ERT2^-WPRE-polyA* cassette was inserted in the first exon of *Col1a2* gene in frame with the translational starting codon between the 5’UTR and the coding sequence (CDS) (**Figure S5D and S5E**). To confirm that the *Col1a2-Cre^ERT2^* line could lead to specific *Map3k2* gene deletion in the colon stromal cells, we sorted the CD326^+^ IEC, CD45^+^ leukocytes, CD31^+^ endothelial cells (CD45^-^CD326^-^CD31^+^), and gp38^+^ intestinal mesenchymal stromal cells (IMSC) (CD45^-^CD31^-^CD326^-^gp38^+^) from the tamoxifen treated *Map3k2^fl/fl^* and *Col1a2-Cre^ERT^:Map3k2^fl/fl^* mouse colon, and performed qRT-PCR for the expression of *Map3k2*. As shown in Figure 6A, we found that *Map3k2* expression was efficiently and specifically ablated only in IMSC from the tamoxifen treated *Col1a2-Cre^ERT2^:Map3k2^fl/fl^* mouse colon but not from the control tamoxifen treated *Map3k2^fl/fl^* mouse colon. Furthermore, *Map3k2* expression was not affected in CD326^+^ IEC, CD45^+^ leukocytes, or CD31^+^ endothelial cells sorted from either the tamoxifen treated control *Map3k2^fl/fl^* or *Col1a2-Cre^ERT2^:Map3k2^fl/fl^* mouse colon (**Figure 6A**). This is consistent with our scRNA-seq data that *Col1a2* is specifically expressed in colon stromal cells (data not shown). Together these data demonstrate that tamoxifen treatment of *Col1a2-Cre^ERT2^:Map3k2^fl/fl^* mice could achieve stromal specific deletion of *Map3k2* in mouse colon.

**Figure 6.**
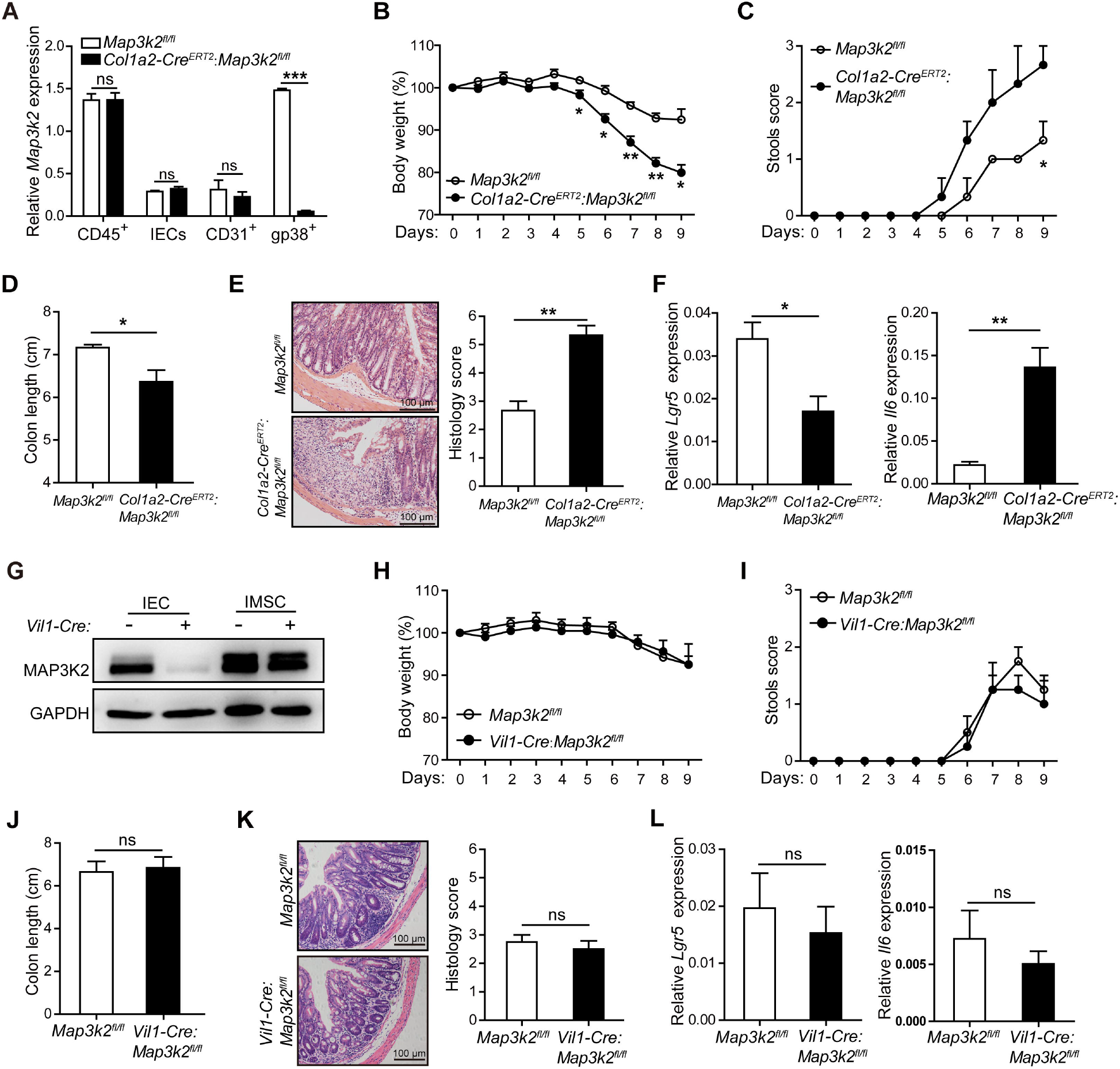
*Map3k2* in Intestinal Stromal Cells Protects Mice from DSS-Induced Colitis *in vivo*. (A) qRT-PCR analysis of *Map3k2* mRNA expression in flow cytometry-sorted CD326^+^CD45^-^ intestinal epithelial cells (IECs), CD45^+^CD326^-^ leukocytes (CD45^+^), CD31^+^CD45^-^CD326^-^ endothelial cells (CD31^+^), and gp38^+^CD45^-^CD31^-^CD326^-^ intestinal mesenchymal stromal cells (gp38^+^) from the colons of tamoxifen treated *Map3k2^fl/fl^* and *Col1a2-Cre^ERT2^:Map3k2^fl/fl^* mice. Data presented are normalized to *Hprt* expression level and are representative of two independent experiments. (B) *Map3k2^fl/fl^* and *Col1a2-Cre ^ERT2^:Map3k2^fl/fl^* co-housed littermate mice (n=3/group, 8 weeks old) were injected intraperitoneally with tamoxifen (TM) every two days (2 mg/mouse/time, 5 times). After 3 weeks from the last time of injection, mice were administered with 2% DSS in drinking water for 7 days followed by regular drinking water for additional 2 days. Body weights were recorded daily. (C) Daily stool scores for mice treated as described in panel (B). (D) Bar graphs show colon length of mice treated as described in panel (B) before sacrifice on day 9. (E) Hematoxylin and eosin (H&E) staining of colon (left panels) and histological assessment of the mucosa (right panels) in mice treated as described in panel (B) sacrificed on day 9. (F) Colonic mRNA expression levels of *Lgr5* (intestinal stem cell marker) and *Il6* in mice treated as described in panel (B) sacrificed on day 9. Data shown are normalized to *Hprt* expression level. (G) Immunoblot analysis of MAP3K2 expression in intestinal epithelial cells (IEC) and intestinal mesenchymal stromal cells (IMSC) isolated from the colons of *Map3k2^fl/fl^* and *Vil1-Cre:Map3k2^fl/fl^* mice. The GAPDH level was determined as loading controls. (H) *Map3k2^fl/fl^* and *Vil1-Cre:Map3k2^fl/fl^* co-housed littermate mice (n=4 pairs) were administered with 2% DSS in drinking water for 7 days followed by regular drinking water for additional 2 days. Body weights were recorded daily. Shown is a representative graph of two independent experiments. (I) Daily stool scores for mice treated as described in panel (H). (J) Bar graphs show colon length of mice treated as described in panel (H) before sacrifice on day 9. (K) Hematoxylin and eosin (H&E) staining of colon (left panels) and histological assessment of the mucosa (right panels) in mice treated as described in panel (H) before sacrifice on day 9. (L) Colonic mRNA expression levels of *Lgr5* and *Il6* in mice treated as described in panel (H) before sacrifice on day 9. Data shown are normalized to *Hprt* expression level. Error bars indicate mean ± SEM (*p<0.05, **p<0.01 ***p<0.001 by unpaired Student’s t test).

We next performed DSS-induced colitis experiments with co-housed littermates of *Map3k2^fl/fl^* and *Col1a2-Cre^ERT2^:Map3k2^fl/fl^* mice after tamoxifen treatment. Within 5-6 days of DSS intake, we observed that *Col1a2-Cre^ERT2^:Map3k2^fl/fl^* mice suffered greater loss of body weight (**Figure 6B**) and worse diarrhea (**Figure 6C**) than did *Map3k2^fl/fl^* mice. *Col1a2-Cre^ERT2^:Map3k2^fl/fl^* mice also exhibited significantly shorter colon length (**Figure 6D**) as well as more severe epithelial damage and greater leukocyte infiltration than were observed in *Map3k2^fl/fl^* mice on day 9 after DSS treatment (**Figure 6E**). Furthermore, we observed reduced *Lgr5* expression and increased *Il6* expression in *Col1a2-Cre^ERT2^:Map3k2^fl/fl^* mouse colon as compared to that in *Map3k2^fl/fl^* mouse colon suggesting more stem cell loss and augmented inflammation (**Figure 6F**). These data together reveal that *Map3k2* plays a stromal specific role *in vivo* in host protection against acute intestinal tissue damage.

To further rule out the potential role of *Map3k2* in intestinal epithelial cells in protection of mice from DSS-induced colitis, we also generated epithelial specific *Map3k2* conditional KO mice, by crossing the *Map3k2^fl/fl^* line with *Vil1-Cre* mice, an intestinal epithelial cell specific *Cre* line (Madison et al., 2002). The expression of *Map3k2* was efficiently ablated in purified IEC but not IMSC from the *Vil1-Cre:Map3k2^fl/fl^* mice colon (**Figure 6G**). Using a similar mouse colitis model described above, we found that *Vil1-Cre:Map3k2^fl/fl^* mice developed similar degree of DSS-induced colitis as that of the control *Map3k2^fl/fl^* mice as measured by body weight loss (**Figure 6H**), diarrhea scores (**Figure 6I**), colon length change (**Figure 6J**), and tissue damage severity (**Figure 6K**).

There was also no significant difference in the stem cell loss or inflammation level in the colons of control *Map3k2^fl/fl^* mice and *Vil1-Cre:Map3k2^fl/fl^* mice as measured by *Lgr5* and *Il6* expression (**Figure 6L**). Finally, we also generated *CD4-Cre:Map3k2^fl/fl^* mice to rule out T cell specific role of *Map3k2* in protection of mice from DSS induced colitis, and found no significant difference in DSS-induced colitis between the *Map3k2^fl/fl^* mice and *CD4-Cre:Map3k2^fl/fl^* mice (**Data not shown**).

## Discussion

Mesenchymal stromal cells are a heterogeneous population that exerts a range of structural and regulatory functions in the intestine. Multiple subsets of stromal cells have recently been shown to provide critical support for the ISC compartment (Kinchen et al., 2018), but it has remained unclear whether a specific lineage of niche-specific stromal cells regulates ISC proliferation and survival in response to acute tissue damage. In the current study, we identify a novel intestinal population of CD90^mid^CD81^+^CD34^+^ mesenchymal stromal cells that are major source of the Wnt agonist *Rspo1in* colon. Anatomically, these *Map3k2*-regulated intestinal stromal cells (MRISC) are located near the bottom of the colonic crypts, where we propose they are likely to form a unique intestinal stem cell niche. The ability of MRISC to augment *Rspo1* expression following colonic injury reflects a likely critical role in sustaining Wnt signaling in intestinal stem cells, leading to rapid epithelial regeneration and repair.

Similar to their roles in other organs and tissues, mesenchymal cells in the intestine are key regulators of epithelial homeostasis, matrix remodeling, immunity, and inflammation (Owens and Simmons, 2013; Pinchuk et al., 2010; Powell et al., 2011). While mesenchymal stromal cells in the gut express a range of common marker genes including *Pdpn, Vim, Col1a, Col1b, Col6a*, increasing evidence suggests that there is extensive heterogeneity within this compartment (Kinchen et al., 2018). However, at present there are no known phenotypic markers that allow unique stromal cell subsets to be distinguished within this heterogeneous group. In the current study, our analyses of *Map3k2* function in experimental colitis revealed that a specific subset of stromal cells critically depends on this enzyme to mediate damage-induced *Rspo1* expression. Further investigation confirmed that this novel population displayed moderate expression of CD90, which subsequently analysis of this subset by multi-parameter flow cytometry (although lymphatic endothelial cells [LEC] located within the same sub-crypt region express 10-fold higher levels of CD90 that can complicate immunostaining approaches). Specific surface markers that define MRISC were identified using scRNA-seq, which revealed that these stromal cells also express CD81 and CD34 but lack CD31 and CD138. When used in combination, these markers facilitated MRISC visualization without the need for CD90 staining, which allowed subsequent confirmation that MRISC cells are located near to *Lgr5*^+^intestinal stem cells within the sub-crypt region. Indeed, MRISC also appeared to interact with adjacent LECs and local immune cells, suggesting that this lineage may indeed organize multiple cell types into a specific stem cell niche in the gut.

Intriguingly, while our data demonstrate that MRISC are critically important for regeneration and repair of the damaged intestine, these cells, or at least the *Map3k2-Rspo1* axis in these cells, may not be essential for normal gut maintenance or homeostasis. These findings contrast with the recent identification of telocytes and *Gli1*-expressing stromal cells that appear critical for steady-state maintenance of the intestine (Degirmenci et al., 2018; Shoshkes-Carmel et al., 2018).

Indeed, while telocytes are *Foxl1^+^* stromal cells that populate the sub-epithelium from stomach to colon, we observed that MRISC are *Foxl1^-^* cells with distinct morphology and localization characteristics. Similarly, while *Gli1*-expressing stromal cells (including *Foxl1^+^* telocytes) are essential sources of Wnt in the intestine, it is unknown whether these cells are involved in driving colonic *Rspo1* expression and stem cell proliferation in the damaged gut as was demonstrated for MRISC. Based on these differences in gene expression, cell morphology, distribution, and unique responsiveness to tissue damage, we propose that MRISC are a novel population of intestinal stromal cells with important niche functions distinct from either telocytes or Gli1-expressing stromal subsets.

Wnt and BMP ligands form a gradient of antagonistic signals in the gut that are known to regulate epithelial renewal and crypt-villi maintenance (Angerer et al., 2016). While the CD81^+^CD90^mid^ MRISC identified in our study might be predicted to produce *Rspo1* and maintain a steady level of Wnt signaling in the intestinal stem cell niche, little is known about how Wnt signals are regulated following acute intestinal damage. In addition to *Rspo1*, MRISC were also observed to express *Rspo3* and Wnt ligands *Wnt2b, Wnt5a* and the BMP pathway inhibitors Gremlin1, Gremlin2, further indicating that MRISC may also have a potential role in supporting ISC niche.

Gut microbiome has been shown to be critical for intestinal homeostasis and tissue regeneration (Elinav et al., 2011). It is possible that the *Map3k2* mediated protection of gut damage might be affected by commensal bacteria or *Map3k2* deficient mice may have a disorder in gut microbiome leading to more severe colitis. However, all our mice were co-housed and treated with DSS together. In addition, we also delivered antibiotics cocktail (Ampicillin, Vancomycin, Neomycin, Metronidazole, and Gentamycin) to either WT or *Map3k2* deficient mice before and during colitis induction with DSS, and observed similar results in co-housed mice as those nontreated mice (**Data Not Shown**). Therefore, the host microbiome is unlikely the major factor for the *Map3k2* deficiency caused severe gut damage.

In summary, our study reveals that a novel population of *Map3k2*-regulated intestinal stromal cells (MRISC) is crucial for induction of *Rspo1* and Wnt signaling which promote rapid stem cell proliferation in response to acute intestinal damage. The sub-crypt location of MRISC likely reflects a requirement for direct communication with ISC following tissue injury, although it is currently unclear whether MRISC expression of additional mediators such as IL-6 and IL-33 reflects an ability to interact with other nearby cell types e.g. ILCs, macrophages, and neutrophils. Deeper investigation of the roles played by MRISC in restraining intestinal inflammation and immunopathology will shed important new light on a range of major pathological conditions. In particular, these studies may provide new rationales for use of MAPK inhibitors or Wnt modulators for the treatment of IBD and colitis-associated colorectal cancer.

## Acknowledgements

We would like to thank Drs. Yuan Zhuang, Florent Ginhoux, Jinke Cheng, Yu Li, and Qijun Wang for reading of our manuscript and valuable suggestions. We also want to thank Drs. Zizhen Kang, Honglin Wang for sharing mice. This work was supported in part by grants from the National Natural Science Foundation of China (31470845, 81430033 and 81871269), Shanghai Science and Technology Commission (13JC1404700), Scientific Research Project of Shanghai Municipal Commission of Health and Family Planning (201640137) and the Interdisciplinary Program of Shanghai JiaoTong University (YG2014MS77).

