## Supplementary figures and images for "*Map3k2*-Regulated Intestinal Stromal Cells (MRISC) Define a Distinct Sub-cryptic Stem Cell Niche for Damage Induced Wnt Agonist R-spondin1 Production"

### Supplemental Figure 1

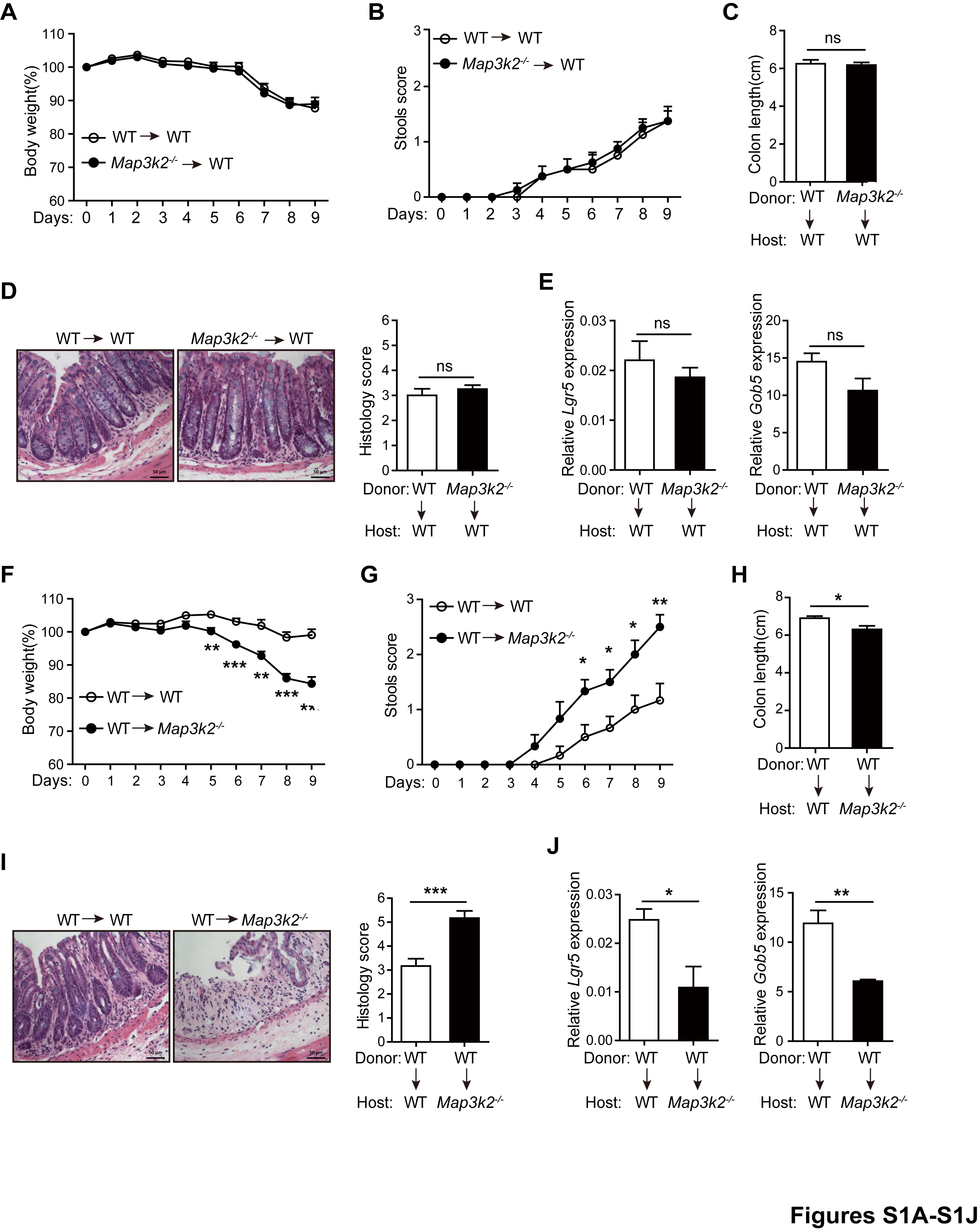

### Supplemental Figure 2

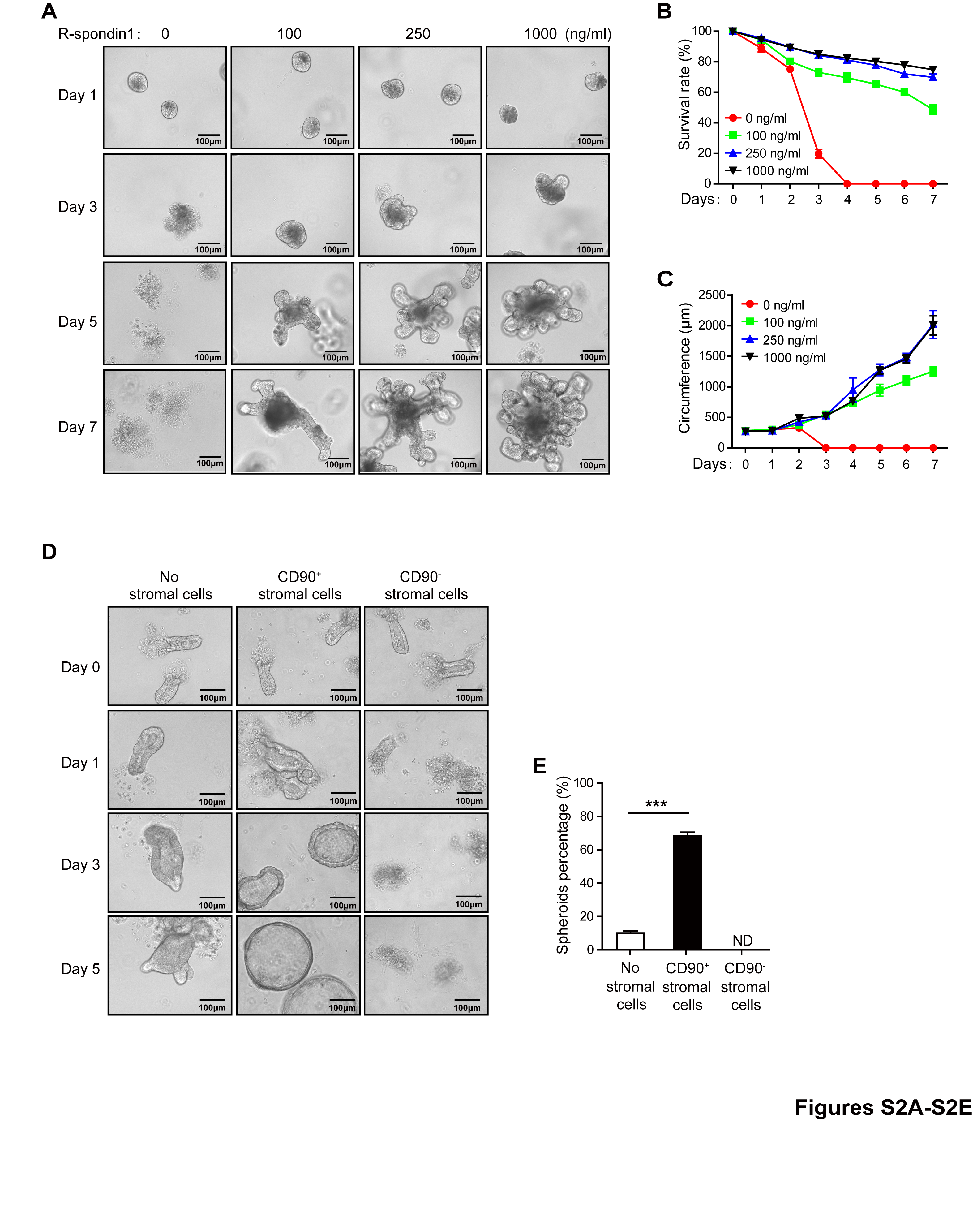

### Supplemental Figure 3

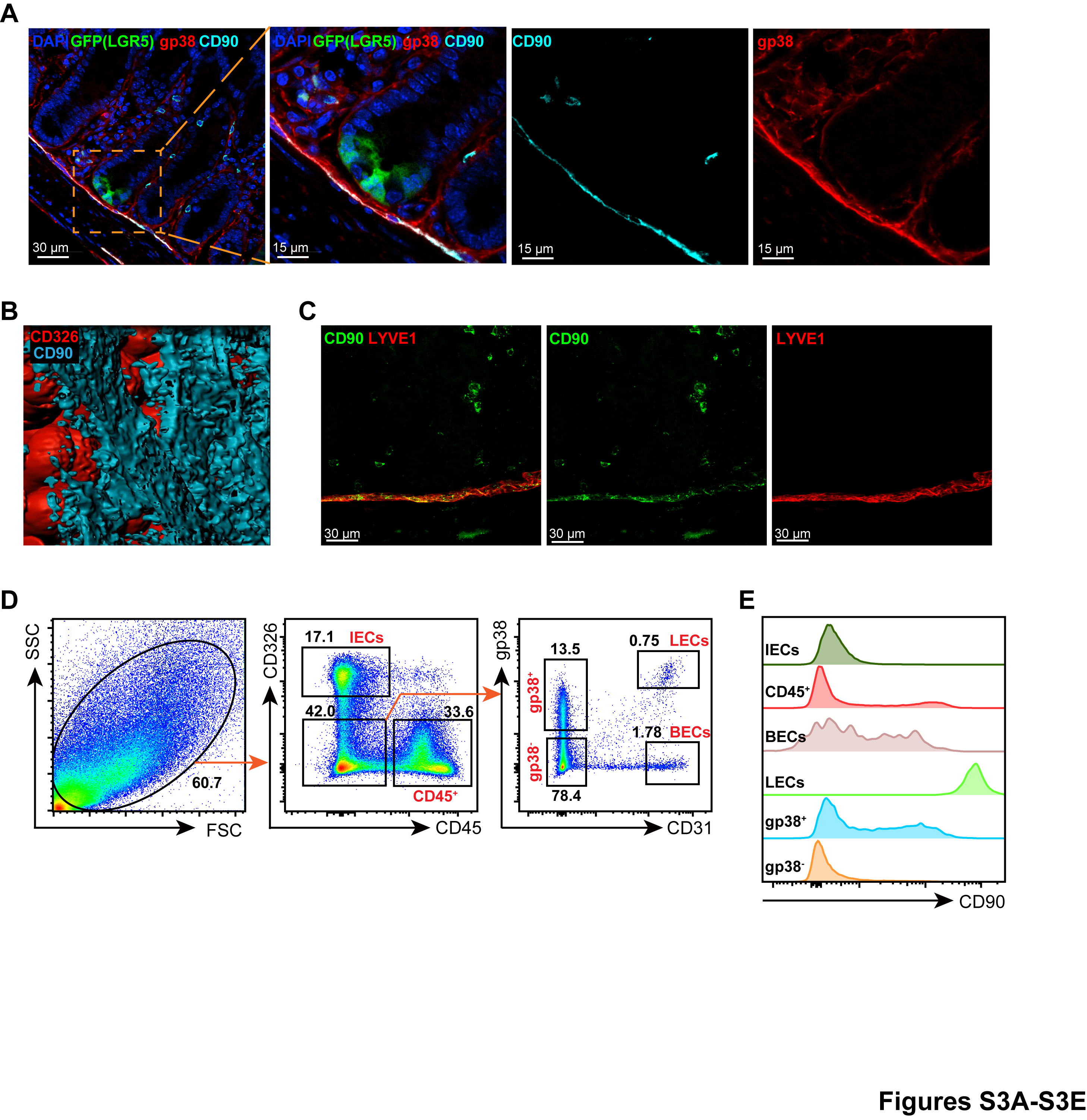

### Supplemental Figure 4

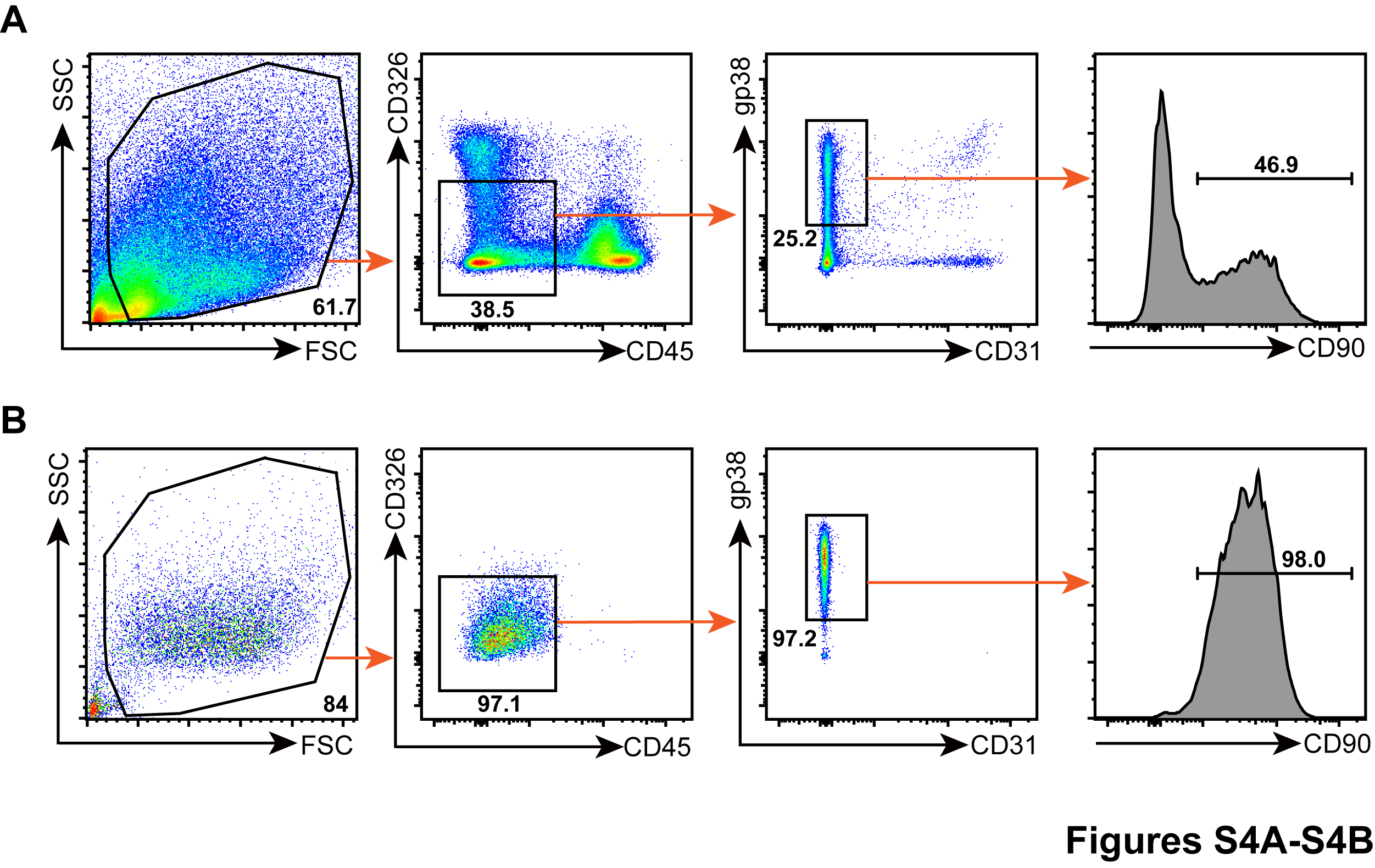

### Supplemental Figure 5

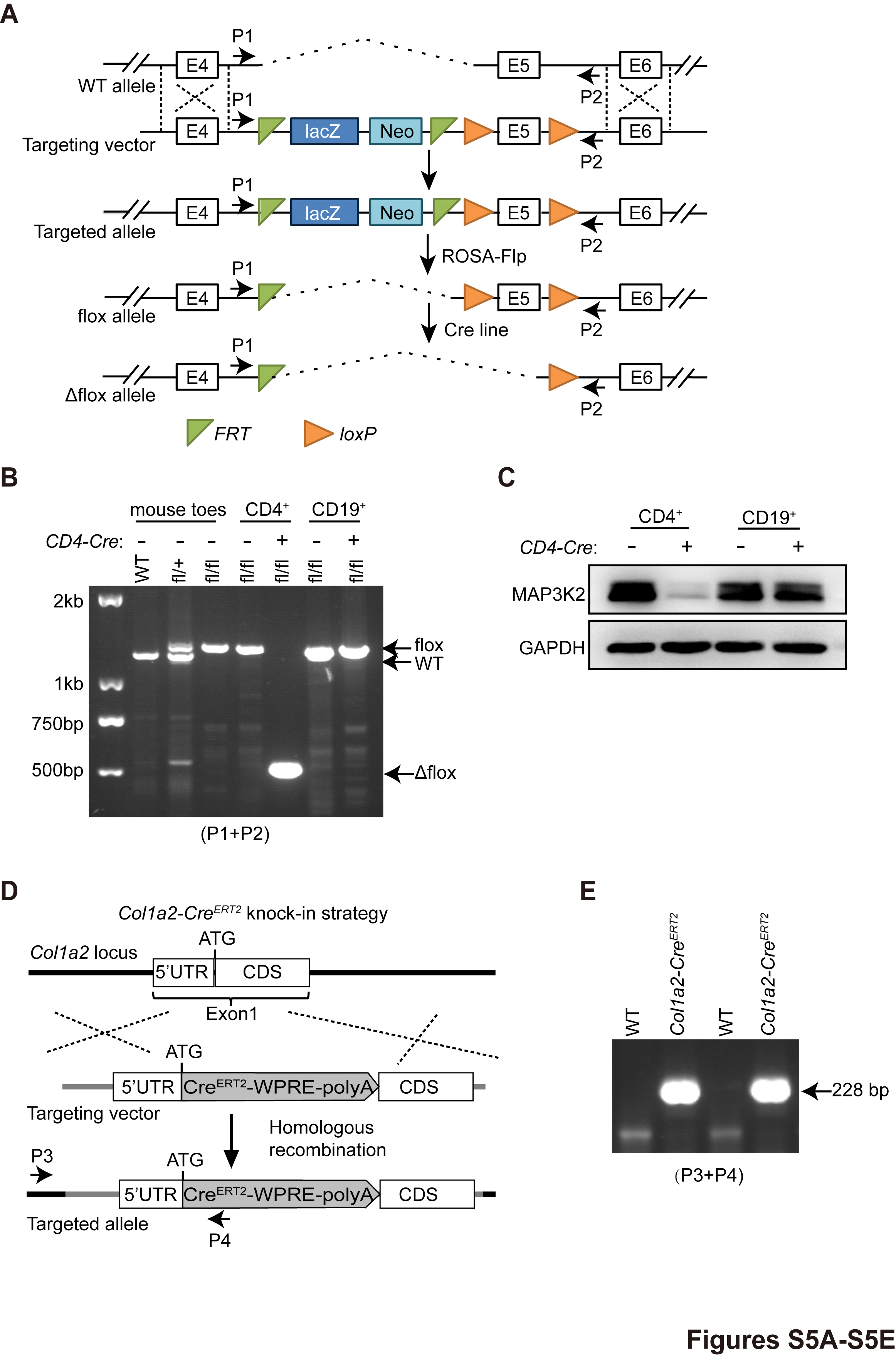
