## Supplemental Figure legends for "*Map3k2*-Regulated Intestinal Stromal Cells (MRISC) Define a Distinct Sub-cryptic Stem Cell Niche for Damage Induced Wnt Agonist R-spondin1 Production"

**Figure S1. Non-hematopoietic *Map3K2* Is Required for Protection against DSS Colitis. Related to Figure 1 and Figure 2**

(A) Daily body weight scores of irradiated WT mice transplanted with bone marrow cells from either WT or *Map3k2^-/-^* donors via *i.v.* administration (n=8 mice/group). Adoptive transfers were performed 2 months before treating the mice with 2% DSS in drinking water for 7 days (followed by provision of regular drinking water for 2 days thereafter).

(B) Daily stool scores for mice treated as described in panel (A).

(C) Colon length of mice treated as described in panel (A) before sacrifice on day 9.

(D) Representative H&E staining (left) and histological assessment of colonic mucosa (right) in mice treated as described in panel (A) before sacrifice on day 9.

(E) Colonic mRNA expression levels of *Lgr5* (intestinal stem cell marker) and *Gob5* (goblet cell marker) in mice treated as described in panel (A) before sacrifice on day 9. Data shown are normalized to *Hprt* expression level.

(F) Daily body weight scores after transplant of bone marrow cells from WT animals into lethally-irradiated WT recipients (WT to WT) or *Map3k2^-/-^* recipients (WT to *Map3k2^-/-^*) 2 months prior to administration of 2% DSS in drinking water for 7 days (followed by provision of regular drinking water for a further 2 days) (n=6 mice/group).

(G) Daily stool scores for mice treated as described in panel (F).

(H) Colon length in mice treated as described in panel (F) before sacrifice on day 9.

(I) Representative H&E staining (left) and histopathology scoring of colon sections (right) in mice treated as described in panel (F) before sacrifice on day 9.

(J) Colonic mRNA expression levels of *Lgr5* (intestinal stem cell marker) and *Gob5* (goblet cell marker) in mice treated as described in panel (F) before sacrifice on day 9. Data shown are normalized to *Hprt* expression level.

Error bars indicate mean ± SEM (*p<0.05, **p<0.01, ***p<0.001 by unpaired Student’s t test).

**Figure S2.** **R-spondin1 Is Required for Intestinal Stem Cell Growth and Survival *In Vitro*. Related to Figure 3 and Figure 4**

(A) Bright-field images of small intestinal crypts isolated from WT mice and cultured in matrigel for 1-7 days in the presence of EGF and Noggin with or without the indicated concentrations of R-spondin1.

(B) Statistical analysis of average organoid number/survival derived from mice treated as described in panel (A) (n=3/group).

(C) Statistical analysis of average organoid circumferences derived from mice treated as described in panel (A).

(D) Bright-field images of colonic crypts isolated from WT mice and cultured in matrigel for 1-5 days either alone in the presence of EGF, Noggin, Wnt3a and R-spondin1 or together with WT CD90^+^ stromal cells (CD45^-^, CD326^-^, CD31^-^, gp38^+^, CD90^+^) or CD90^-^ stromal cells (CD45^-^, CD326^-^, CD31^-^, gp38^+^, CD90^-^) in the presence of EGF, Noggin and Wnt3a.

(E) Percentage of spheroids formed among colonoids derived from the cultures described in panel (D) before assessment on day 5 (n=3/group). Error bars indicate mean ± SEM (***p<0.001 by unpaired Student’s t test).

**Figure S3. CD90 is Highly Expressed in LECs and Moderately Expressed in Stromal Cells. Related to Figure 4.**

(A) Immunofluorescence images of swiss roll sections prepared from *Lgr5*-EGFP mouse colon and stained with anti-CD90 (cyan), gp38 (red) and DAPI (blue) for subsequent analysis using a Leica SP8 microscope with 63x oil objective lens. Pictures on right show magnified images of the sub-crypt area highlighted by the dotted square (representative of two independent experiments).

(B) Whole mount sections (3mm x 3mm) of WT mouse colon after clearing and staining with anti-CD90 (cyan) and CD326 (red) for analysis using a Leica SP8 microscope with 20x objective lens.

(C) Images of WT mouse colon processed into swiss roll sections and stained for CD90 (green) and LYVE1 (red) then visualized using a Leica SP8 microscope with 63x oil objective lens.

(D) Representative flow cytometry staining plots and percentage of IEC (CD326^+^), CD45^+^ leukocytes, LECs (CD45^-^, CD326^-^, CD31^+^, gp38^+^), BECs (CD45^-^, CD326^-^, CD31^+^, gp38^-^), gp38^+^ stromal cells (CD45^-^, CD326^-^, CD31^-^, gp38^+^) and gp38^-^ cells (CD45^-^, CD326^-^, CD31^-^, gp38^-^) derived from WT colonic lamina propria. Plots shown are representative of three independent experiments.

(E) Flow cytometric quantification of CD90 expression in the indicated cell types sorted as described in panel (D). Plots shown are representative of three independent experiments.

**Figure S4. Flow Cytometry Sorting Strategy and Purity of CD90^mid^ Stromal Cells Isolated for scRNA-seq. Related to Figure 5**

(A) Representative flow cytometry staining plots showing the percentages of colonic CD90^mid^ stromal cells (CD45^-^CD326^-^CD31^-^gp38^+^CD90^mid^) present in WT colonic lamina propria before sorting.

(B) Representative flow cytometry staining plots showing the percentages of colonic CD90^mid^ stromal cells (CD45^-^CD326^-^CD31^-^gp38^+^CD90^mid^) present in WT colonic lamina propria after sorting.

**Figure S5. Generation of *Map3k2* Conditional Knockout Mice and *Col1a2-Cre^ERT2^* Mice. Related to Figure 6**

(A) A diagram showing the *Map3k2* gene locus (WT allele), the targeting vector, the targeted allele (Targeted allele), the loxP sequence-targeted allele with the Neo gene deleted by ROSA-Flp (flox allele), and the conditionally deleted allele by a Cre line (△flox allele) as indicated. E4, E5 and E6 indicate the exon 4, exon 5 and exon 6 of *Map3k2* gene, respectively. P1 (GATTCTTTTCTGGGTGTTTGCTAGC) and P2 (CTGCAACTCATAAACCCTCACATCC) are primers used for PCR reactions to distinguish the WT, flox and △flox of *Map3k2* alleles, and their relative locations in the *Map3k2* gene locus are shown in the diagram as indicated.

(B) PCR genotyping of the *Map3k2* conditionally targeted mice with P1 and P2 primers. DNAs were prepared from mouse toes of WT, *Map3k2^fl/+^* and *Map3k2^fl/fl^* mice*,* or from CD4^+^ T cells and CD19^+^ B cells sorted from the spleen of *Map3k2^fl/fl^* and *CD4-Cre*:*Map3k2^fl/fl^* mice. WT, flox, and △flox alleles are indicated by the arrows.

(C) Immunoblot analysis of MAP3K2 expression in CD4^+^ T cells and CD19^+^ B cells sorted from the spleen of *Map3k2^fl/fl^* and *CD4-Cre*:*Map3k2^fl/fl^* mice. The GAPDH level was determined as a loading control.

(D) A diagram illustrating the *Col1a2* gene locus of exon 1 containing the 5’ untranslated region (5’UTR), translation starting codon ATG, and the immediate downstream coding sequence (CDS); the targeting vector which contains a *CreER-WPRE-polyA* cassette inserted in-frame with the ATG codon of the *Col1a2* gene in exon 1 just downstream of 5’UTR; and the targeted allele (*Col1a2-Cre^ERT2^*) in which the *CreER-WPRE-polyA* cassette is inserted in the exon 1 as indicated. P3 (ATTATTTTAGCACCACGGCAGC) and P4 (TGCGAACCTCATCACTCGTTG) are primers used for PCR reactions to distinguish the WT and *Col1a2-Cre^ERT2^* alleles, and their relative locations are shown in the diagram.

(E) PCR genotyping of the *Col1a2-Cre^ERT2^* mice with P3 and P4 primers. DNAs were prepared from mouse toes isolated from WT and *Col1a2-Cre^ERT2^* mice as indicated.
