## Supplementary material for "*Map3k2*-Regulated Intestinal Stromal Cells (MRISC) Define a Distinct Sub-cryptic Stem Cell Niche for Damage Induced Wnt Agonist R-spondin1 Production": Methods

**CONTACT FOR REAGENT AND RESOURCE SHARING**

**EXPERIMENTAL MODEL AND SUBJECT DETAILS**

**Mice**

*Map3k2^-/-^* mice have been described previously (Guo, Clydesdale et al. 2002) and were bred with C57BL/6 mice for more than 10 generations prior to use in the experiments. *Map3k2* floxed mouse line and *Col1a2-Cre^ERT2^* mouse line were generated by Shanghai Model Organisms Center, Inc. (SMOC). The genotyping PCR primer sequences are shown in Table S1. *Lgr5*-EGFP-IRES-*Cre^ERT2^* (*Lgr5*-EGFP), *CD4-Cre*, *Vil1-Cre,* and *Pdgfra-Cre^ERTM^* mice were obtained from the Jackson Laboratory. All mice were bred and maintained at accredited animal facilities under specific pathogen-free conditions in individually ventilated cages on a strict 12h day/night cycle with a regular chow diet. Unless otherwise indicated, 10-16 weeks old and sex-matched mice were used in all assays. All animal experiments were performed according to local guidelines for the use and care of laboratory animals as provided by Shanghai Jiao Tong University School of Medicine Institutional Animal Care and Use Committees (IACUC).

**DSS-Induced Colitis and Determination of Stool Scores**

Mice aged 10-16 weeks and gender-matched were administered 2% DSS (M.W.= 36,000–50,000 Da; MP Biomedical) in sterile drinking water over a period of 7 days, followed by provision of normal drinking water for a further 2 days. Mice were weighed daily and scored for stool consistency. According to protocol, mice were sacrificed if they lost more than 30% of initial body weight during colitis induction. Stool scores were determined as follows: 0, well-formed pellets; 1, soft but still formed; 2, semi-formed stools that adhered to the anus; 3, liquid stools that adhered to the anus.

**Bone Marrow Chimeras**

Bone marrow samples were collected from the shinbone and thighbone of donor mice (CD45.1) and re-suspended in PBS at a concentration of 1×10^7^/ml before a 200μl volume was injected intravenously into lethally-irradiated recipient mice (CD45.2). After 7 weeks, bone marrow reconstitution was verified by blood cell staining with FITC anti-mouse CD45.1 (eBioscience) and PerCP-Cy5.5 anti-mouse CD45.2 antibody (eBioscience). After 8 weeks recovery, the mice were subjected to the DSS-induced colitis protocol described above.

**Isolation and Culture of Mouse Colonic Stromal Cells**

Mouse colons were surgically excised, flushed with PBS, then opened longitudinally and cut into ~0.5cm length pieces. The colon tissue was then washed further and twice incubated in HBSS containing 2% FBS, 1mM EDTA and 1mM DTT, at 37℃ with constant shaking at 200rpm for 20min to release epithelial cells. The mucosal tissues were then washed with DMEM containing 10% FBS and digested in DMEM containing 10% FBS, collagenase type Ⅷ (50units/ml, Sigma) and DNase Ⅰ (50 units/ml, Sigma) at 37℃ for 50min. After digestion, the remaining tissue fragments were collected into a 50ml tube and vortexed vigorously for 30s and passed through a 100μm cell strainer. The resultant cell suspension was then centrifuged at 1500rpm, 4℃ for 5min before discarding the supernatant and re-suspending the cell pellets in DMEM containing 15% FBS, 100U/ml penicillin, and 100μg/ml streptomycin (Invitrogen). After 24h culture, non-adherent cells were removed prior to addition of 10ml fresh DMEM supplemented with 15% FBS, 100 U/ml penicillin, and 100μg/ml streptomycin (Invitrogen). After 4 days culture, stromal cells were purified by FACS sorting based on staining with anti-CD90 (BioLegend). The percentage of CD90^+^ cells was routinely assessed as >90%.

**Intestinal Organoid Culture**

Colon and small intestine were gently removed from the mouse abdominal cavity, opened longitudinally, and then cut into 1cm length pieces. The intestinal fragments were washed 4 times in a 20ml volume of ice-cold DPBS in a 50ml tube with gentle shaking, followed by 30min incubation in 2mM EDTA/DPBS at 4℃ with further gentle shaking. Crypts were released by shaking the tube for 20s in 5ml ice-cold dissociation buffer (54.9mM D-sorbitol and 43.4mM sucrose in DPBS), then passed through a 100μm filter and collected by centrifugation at 150g for 5min, 4℃. The isolated crypts were then mixed with growth factor-reduced Matrigel (Corning) and placed at the center of each chamber in a 24-well plate (200 crypts/50μl Matrigel). Following polymerization of the Matrigel for 15min at 37℃, each well was supplemented with 500μl advanced DMEM/F12 medium (Life Technologies) containing 200ng/ml EGF (Life Technologies), 100ng/ml Noggin (PeproTech) and 1µg/ml R-spondin1 (PeproTech). For colonoid cultures, an additional 100ng/ml Wnt3a (R&D Systems) was added as required. The crypts were incubated at 37℃ with 5% CO_2_ for 5 days. The frequencies and circumferences of colonic organoids were recorded daily. For cell type analyses, colonoids were collected at day 5 and treated with TRIzol reagent for total RNA isolation.

For co-culture experiments, a total of 200 crypts were mixed with 1x10^5^ flow-sorted colonic CD90^+^ stromal cells (CD45^-^CD326^-^CD31^-^gp38^+^CD90^+^) or CD90^-^ stromal cells (CD45^-^CD326^-^CD31^-^gp38^+^CD90^-^) and embedded in matrigel (Corning). Following polymerization of the Matrigel (15min at 37℃), cultures were supplemented with 500μl Advanced DMEM/F12 (Life Technologies) containing EGF (200ng/ml, Life Technologies), Noggin (100ng/ml, PeproTech) and Wnt3a (100ng/ml, R&D). Number and circumference of colonic organoids and spheroids were recorded daily.

***In vivo* R-spondin1 Administration**

Recombinant human R-spondin1 (Peprotech) was dissolved in ddH2O to a stock concentration of 500μg/ml and then stored at -80℃. For *in vivo* administration experiments, the recR-spondin1 stock was diluted to 25μg/ml in saline immediately before use. *Map3k2^-/-^* mice aged 10-12 weeks and gender-matched were randomly divided into two cohorts for administration of either 5µg recR-spondin1 per mouse (i.p.) or a matching volume of saline only (i.p.). After 24 hours, both cohorts were subjected to DSS-induced colitis as described above. For the duration of colitis induction, recR-spondin1 or saline control were administrated daily via i.p. injection.

**NAC Treatment *In Vivo***

WT mice aged 8-10 weeks were administered 2% DSS or regular drinking water (untreated) for 2 days. *N-*acetyl-L-cysteine (NAC) (Sigma) was injected at a daily dose of 27.5mg/kg/day (i.p.). Control mice received PBS injection only (i.p.). Colon tissue was excised and used to prepare total RNA for qRT-PCR as described above.

**Tamoxifen Induced *Map3k2* deletion**

*Map3k2^fl/fl^* mice and *Col1a2-Cre ^ERT2^Map3k2^fl/fl^* co-housed littermates (8 weeks old) were injected intraperitoneally with tamoxifen (TM) every two days (2 mg/mouse/time, 5 times). After 3 weeks from the last time of injection, *Map3k2* deletion was confirmed by PCR and immunoblotting in flow cytometry sorted gp38^+^ IMSC.

**METHOD DETAILS**

**RNA Extraction and Quantitative Real-Time PCR (qRT-PCR)**

Mice were subjected to various experimental procedures as described in the text then sacrificed and the colon tissue resected and frozen using liquid nitrogen. The frozen tissues were disaggregated using a motorized homogenizer followed by RNA extraction using the RNeasy Mini Kit (QIAGEN). Primary colon-derived mesenchymal stromal cells and FACS-sorted lamina propria populations were lysed with 1ml TRIzol® Reagent (Invitrogen) and total RNA was isolated according to standard protocols. Total RNA was then immediately reverse-transcribed into cDNA using the PrimeScript™RT reagent Kit (Perfect Real Time) (Takara). qRT-PCR was subsequently performed using SYBR Green Real-time PCR Master Mix (Takara) together with a ViiA7 Real-Time PCR System (Applied Biosystems). All primers were purchased from Sangon Biotech. The qRT-PCR primer sequences are shown in Table S2.

**Gene Expression Profiling of Colon Tissues using Next-Generation Sequencing**

At 10-12 weeks after birth, age/gender-matched WT and *Map3k2^-/-^* mice were administered 2% DSS (MP Biomedical) in drinking water for either 0 or 24h. Colon tissues were collected and total RNA was isolated using RNeasy Mini Kits for quantification using Qubit 3. cDNA libraries were then generated using the [TruSeq Stranded mRNA Library Prep Kit](https://www.illumina.com/products/by-type/sequencing-kits/library-prep-kits/truseq-stranded-mrna.html) (Illumina) and sequenced on a NextSeq500 (Illumina) with PE75. Sequences obtained from the RNA-seq pipeline were aligned against the *Mus musculus* genome using Bowtie 2 (Langmead and Salzberg 2012). HTSeq-count (Anders, Pyl et al. 2015) was used to enumerate transcripts associated with each gene, and a counts matrix containing the number of counts for each gene across different samples and stimulations was obtained. The counts were normalized based on sample size. Genes that were differentially expressed in the untreated and DSS-challenged conditions were then compared between WT and *Map3k2^-/-^* mice. A list of genes that showed statistically significant regulation in opposite directions in the WT and MEKK2 knockout was obtained and analyzed for biological pathway enrichment and Transcription Factor Regulation Enrichment in GSEA. For all comparisons, an FDR-adjusted p-value of 0.05 was used as the threshold for statistical significance (with Benjamini-Hochberg test for multiple testing correction).

**Flow Cytometry Analysis and Fluorescence Activated Cell Sorting (FACS)**

Single cell suspensions were stained with fluorophore-conjugated anti-mouse antibodies at the recommended dilutions for 30min at 4℃ in the dark. Cells were then washed once with 1×PBS containing 2% FBS and re-suspended in 1×PBS containing 2% FBS for flow cytometry or cell sorting. The antibodies used in this study were: APC-cy7 anti-mouse Podoplanin PerCP-Cy5.5 anti-mouse CD81, PE-cy7 anti-mouse CD138, BV421 anti-mouse CD34 from BioLegend. APC anti-mouse CD326, and PE anti-mouse CD31 were from eBioscience. FITC anti-mouse CD45, BV605 anti-mouse CD90.2 were from BD Biosciences. For flow cytometry, stained cells were analyzed using a Fortessa X20 instrument with FlowJo software (BD Biosciences). For live sorting experiments, antibody-stained cells were immediately sorted into DMEM containing 20% FBS using a BD FACS Aria III (BD Biosciences).

**Chromatin Immunoprecipitation (ChIP) Assay**

Colonic stromal cells were crosslinked for 20min at RT using formaldehyde (1% final concentration) before quenching the reaction for 5min at RT with glycine to achieve a final concentration of 0.125M. The cells were then washed twice with 5ml ice-cold PBS and re-suspended in lysis buffer (1% Triton X-100, 500mM NaCl, 1mM EDTA, 0.1% Na-Deoxycholate, 0.1% SDS, 1mM PMSF, 50mM HEPES, pH 7.5) and placed on ice for 20min. Using an ice bath to maintain low temperature, the cells were sonicated with a BRANSON DIGITAL SONIFIER 250 for 10min at 50% output using a 5s on/off cycle. Next, a 1.2ml volume of sonicated chromatin suspension was incubated with 2μg anti-KLF2 antibody (Merck Millipore) or 2μg Rabbit IgG control mAb DA1E (CST) for 12h at 4℃ under constant rocking. After incubation, 30μl protein A/G agarose beads were added to each tube for 3h at 4℃ with gentle rocking. The beads were then washed three times using 1ml high salt buffer (1% Triton X-100, 500mM NaCl, 1mM EDTA, 0.1% Na-Deoxycholate, 50mM HEPES, pH 7.5), followed by one wash with 1ml low salt buffer (0.5% NP-40, 250mM LiCl, 1mM EDTA, 0.5% Na-Deoxycholate, 10mM TrisCl, pH8.0), and a final wash with 1ml TE buffer (1mM EDTA, 10mM TrisCl, pH 8.0). Chromatin was then eluted by incubation of the beads at 65℃ for 4h in Elution and De-crosslinking buffer (1% SDS, 10mM EDTA, 50mM TrisCl, pH 8.0) prior to purifying using a QIAquick PCR Purification Kit (QIAGEN) according to the manufacturer’s instructions. qRT-PCR was performed with SYBR Green Real-time PCR Master Mix (Takara) together with a ViiA7 Real-Time PCR System (Applied Biosystems). Primer sequences were as follows:

Klf binding motif in *Rspo1* promotor-F: TGAGGTTCTCTCCATCCAGTCT

Klf binding motif in *Rspo1* promotor-R: TGCAACGACCTAGGAGAGGT

Negative control (20kb to TSS)-F: AGCCATCCCTCCAGTCTCAT

Negative control (20kb to TSS)-R: GCCTGGTTAATGCATGTGTTGT

**Lentivirus Transfection**

EGFP-tagged KLF2 or EGFP alone were cloned into pLVX-IRES-Puromycin vector (Clontech Laboratories, Inc). shRNA targeting KLF2 or control shRNA were cloned into the pLVX-shRNA lentiviral vector (Clontech Laboratories, Inc). HEK293T cells were cultured in in 10cm dishes and transfected with pLVX-KLF2 or pLVX-shRNAs together with the viral envelope plasmid (pMD2.G) and viral packaging plasmid (pSPAX2). After transfection for 48 and 72h, virus particles were collected, pooled, and filtered through a 0.45μm nitrocellulose membrane (Millipore). WT colonic stromal cells were infected with lentivirus encoding KLF2 and EGFP control or KLF2-shRNA and Control-shRNA in the presence of 10mg/ml polybrene (Sigma). On day 4 after infection, cells were stained with anti-CD90 mAb (BioLegend). CD90^+^EGFP^+^ cells were sorted and total RNA was isolated using TRIzol Reagent (Invitrogen). Total RNA was then immediately reverse-transcribed into cDNA using the PrimeScript™RT reagent Kit (Takara). qRT-PCR was performed with SYBR Green Real-time PCR Master Mix (Takara) together with the ViiA7 Real-Time PCR System (Applied Biosystems).

**Immunofluorescence Microscopy**

Colon was harvested, excised and rolled up before fixation in 1% PFA (BBI life sciences) for 24h at 4℃. After dehydrating in 30% sucrose (Shanghai Huixing) for 1 day, the tissues were frozen in Tissue-Tek O.C.T. compound (Sakura Finetek) and cut into 20µm sections for blocking and staining with PBS containing 0.3% TritonX-100 (BBI life sciences), 1% FBS (Jackson ImmunoResearch), 1% BSA (Shanghai Yeasen Biotechnology) and 1% normal mouse serum (Jackson ImmunoResearch). The anti-mouse antibodies used were as follows: CD34 (MEC14.7), Podoplanin (8.1.1), CD326 (G8.8), CD90 (30-H12) and CD31 (MEC13.3) from BioLegend. Anti-CD81 (Eat2) and the goat anti-hamster secondary antibody were from Invitrogen. Images were acquired on a Leica SP8 confocal laser–scanning microscope and processed using Bitplane Imaris 9.1.2.

**RNAscope**

In situ hybridization was performed using the C Multiplex Fluorescent Detection Kit v2 (ACD Bio) according to the manufacturer’s instructions. Colon was excised, rolled up, and immediately frozen in liquid nitrogen before embedding in Tissue-Tek O.C.T. compound (Sakura Finetek). Sections of 10µm thickness were then prepared for RNAscope using a Mm-*Rspo1* probe (ACD Bio 401991) and bacterial *DapB* probe as a negative control (ACD Bio 310043). DAPI was used as a nuclear counterstain.

**ROS Staining by Flow Cytometry**

Mouse colon cells were stained with surface markers at 4℃ for 30 min then washed with FACS buffer, followed by incubation with 10 mM CM-H2DCFDA (Life Technologies) at 37℃ for 15 min. After incubation, fluorescence levels in stained cells were immediately measured by flow cytometry.

**10x Library Preparation and Sequencing**

Cells isolated from mouse colons were re-suspended in PBS containing 0.04% BSA at a final density of 800 cells per μl. The single-cell suspension was mixed with RT-PCR master mix, Single Cell 3′ Gel Beads and Partitioning Oil first, then loaded onto the Single Cell 3′ Chip according to the manufacturer’s instructions (10X Genomics). Briefly, 15,000 cells were loaded into each reaction and the RNA transcripts from individual cells were uniquely barcoded and reverse-transcribed. cDNA molecules were pre-amplified, fragmented, end repaired, and ligated with adapters as per the manufacturer’s protocol in order to generate a single multiplexed library for sequencing in a single Illumina NextSeq 500 cartridge. All libraries were quantified by Qubit and the size profiles of the pre-amplified cDNA and sequencing libraries were examined using an Agilent 2100 BioAnalyzer.

**QUANTIFICATION AND STATISTICAL ANALYSIS**

**10x Genomics Computational Analysis**

**a) Cellranger Pipeline**

Cell Ranger software suite version 2.1 was obtained from 10x Genomics (https://support.10xgenomics.com/single-cell-gene-expression/software/downloads/2.1). Raw sequencing data were first de-multiplexed using Illumina bcl2fastq software to generate separate paired-end read files for each sample. For murine sample libraries, alignment and transcript quantification were performed using the standard Cell Ranger ‘count’ script against the University of California Santa Cruz (UCSC) mm10 murine genome assembly. UMI counts were summarized using the Ensembl gene annotation GTF file obtained using the UCSC Table Browser tool. Filtered count matrices of mouse data were imported into R for further processing.

**b) Seurat Pipeline**

Filtered expression matrices output from the Cell Ranger pipeline were first filtered to remove barcodes with fewer than 250 unique molecules detected. Initial clustering was performed using the R package ‘Seurat’ (Version 2.3.2) (Butler, Hoffman et al. 2018). Low-expressed genes (detected in fewer than 3 cells) were first removed, then variable genes were annotated by examination of the mean-variance relationship. Variable genes met the following criteria: 0.0125 < mean of non-zero values < 3 AND standard deviation > 0.5. Dimension reduction was then performed using PCA and tSNE. The first 53 principle components were used to generate an initial clustering using the Seurat community detection algorithm to identify the cell cluster with resolution of 0.6. Violin plots, feature plots and heat-maps were also generated using 'Seurat' implemented functions.

**Statistical Analysis**

The information about statistical details and methods is indicated in the figure legends or methods. Statistical analysis was performed using Graphpad Prism 5.0. Data are represented as mean ± SEM and analyzed with the unpaired Student’s t test. P<0.05 was considered significant (*p<0.05; **p<0.01; ***p< 0.001).
